# SingleBehavior Lab: behavioral sequencing and phenotyping with lightweight task specific adaptation

**DOI:** 10.64898/2026.04.15.718696

**Authors:** Almir Aljović, Nina Heinrichs, Fritz Kagerer, Hannah Peedle, Florence M. Bareyre

## Abstract

Understanding behavioral differences between experimental groups and quantifying action structure in behavioral experiments remains challenging. Currently, most approaches rely on pose estimation followed by downstream classification, resulting in assay specific pipelines with substantial annotation requirements. Here we present SingleBehavior Lab (SBL), a framework for modeling behavior across experimental contexts using a standardized graphical interface. SBL leverages spatiotemporal embeddings from large video foundation models and combines them with lightweight contrastive adapters, a multi-head attention pooling (MAP) module and a temporal decoder to enable behavior sequencing and task-specific refinement. The framework supports few-shot learning, allowing small models trained on pretrained embeddings to improve action segmentation and classification with limited labeled data, without fine-tuning the underlying video model. In parallel, a large segmentation model with motion-aware memory is used to extract object-centered representations that, together with shared spatiotemporal embeddings, enable unsupervised clustering of behavioral states and analysis of their structure, including cluster prioritization, transition dynamics and attention-based interpretability. Across multiple assays and species, SBL supports identification of group-level differences and rare behaviors, and provides a basis for integrating behavioral representations across experimental contexts.

## Main

Analyzing behavior across experiments, locations and species, and quantifying behavioral differences across these groups remains a central goal in neuroscience and ethology [1]. With the rapid growth of large scale behavioral datasets, the field increasingly requires standardized and adaptable approaches to behavior analysis. Two quantitative directions are particularly prominent in neuroscience experiments. One is understanding the temporal organization and sequencing of actions (here referred to as behavior sequencing), and the other is developing unsupervised, unbiased methods for comparing behavior across groups, enabling insight into circuit perturbations, genetic manipulations, or pharmacological interventions. Recent advances in deep learning and large model training are bringing the field closer to more generalizable behavioral analysis frameworks [2,3].

Currently, the dominant workflow relies on pose-estimation tools, typically trained via transfer learning to extract key points or trajectories, which are then used to classify actions or behavioral states. Markerless pose-estimation systems such as SLEAP and DeepLabCut have made keypoint tracking broadly accessible across species and experimental paradigms, enabling a wide range of downstream behavior analysis pipelines [4,5]. In the context of behavior sequencing, pose-obtained data are often used to train classifiers or sequence models that yield quantitative behavioral phenotypes, supporting both general action classification and more specialized analyses such as social interactions [6–10]. Other methods skip explicit pose estimation, instead operating directly on raw or preprocessed video to predict specific, predefined behaviors. [11,12].

In parallel, unsupervised approaches aim to construct low dimensional behavioral representations and segment behavior into recurring motifs. These include models that decompose continuous motion into meaningful behavioral units and discover action clusters using pose dynamics [13–17], as well as methods that rely on learned video embeddings without explicit pose representations [18–21]. Despite progress along these complementary directions, cross-context adaptation remains a key challenge. Behavior analysis pipelines often require retraining separate pose estimation models and then training downstream classifiers for each new experimental setup, even with standardization of the experimental conditions. Additionally, some of them require specialized equipment like depth cameras or specific cages to enable behavior detection and analysis. Markerless pose-estimation can be further complicated in certain experimental setups, such as those involving miniscopes or complex wiring, where occlusions make reliable keypoint tracking and subsequent action categorization more challenging. Collectively, these limitations suggest that robust behavioral characterization would benefit from representations that transfer across contexts and from adaptation strategies that enable efficient training of task-specific models for behavior sequencing with limited labeled data across diverse experiments.

Recent video foundation models provide a promising path toward transferable behavioral representations by enabling high dimensional spatiotemporal embeddings that can be reused across tasks. VideoPrism, for example, is a general purpose video encoder designed to support diverse downstream video understanding tasks using a single frozen backbone [22]. In the context of behavior analysis, recent evaluations indicate that frozen video foundation models can support animal behavior tasks such as classification, localization, retrieval, and tracking, providing evidence that a single backbone can be repurposed across experimental contexts [23]. A key advantage of these models is that they enable efficient adapter-based learning and small task-specific neural modules trained on top of frozen spatiotemporal embeddings, allowing few-shot adaptations without fine tuning of the backbone.

Based on these considerations, we implemented a process where a foundation video encoder offers a stable spatiotemporal embedding space with lightweight adaptation modules, enabling rapid, few-shot customization for specific behavioral tasks and active model refinement. This approach constitutes a tool that enables the efficient training of lightweight adaptation heads. In addition, we incorporate a foundation segmentation model to support unsupervised behavior discovery through object centric preprocessing [24]. Rather than relying solely on keypoints, this approach uses promptable video segmentation and robust object tracking to extract consistent, object centered clips, enabling embedding and clustering across diverse scenes. In addition we enable downstream analysis tools for cluster inspection and prioritization to facilitate interpretation and enable the detection of rare or anomalous behaviors.

Our toolbox, termed SingleBehavior Lab (SBL), is a framework for cross-context behavioral characterization that uses frozen foundation video embeddings as a shared representational substrate and supports efficient, few-shot behavioral action modeling through contrastive adapter and temporal decoder training. SingleBehavior Lab extracts high dimensional spatiotemporal embeddings from video clips and trains compact contrastive adapters and temporal decoder using augmentation-based objectives to improve action segmentation and classification without fine-tuning the foundation model [25]. This design reduces computational overhead relative to full backbone fine-tuning and enables rapid iteration when labeled data are limited [25–27]. Motivated by single cell analysis paradigms in which integration across conditions is treated as a central challenge, SingleBehavior Lab further supports behaviorome construction across different experiments by integrating shared and context specific behavioral states, conceptually paralleling pseudotime and integration frameworks in single cell genomics [28–30].

We evaluate SingleBehavior Lab across multiple experimental settings and species, including mice, drosophila and marmoset, and demonstrate its utility for behavior sequencing, detection of unique behavioral states, and integration of behavioral states across experiments. We additionally provide a standardized graphical user interface that supports embedding extraction, adapter training, state discovery, object segmentation and tracking as well as downstream sequence and state transition analyses [2,10,24, 31,32].

## Results

### A unified framework for rapid behavior sequencing and unbiased behavioral discovery

We developed SingleBehavior Lab, a framework that enables standardized behavior analysis across experimental contexts by leveraging frozen video foundation model embeddings and lightweight task-specific adaptation modules (**Fig. 1; Extended Data Fig. 1-2**). The framework supports two complementary analysis directions, one is rapid training of behavior classifiers and temporal decoders for behavior sequencing, and the second one is unbiased discovery of behavioral states through unsupervised embedding and clustering. To enable efficient behavior annotation with minimal labeled data, SingleBehavior Lab implements a few-shot learning pipeline built on top of a frozen video foundation encoder (**Fig. 1a; Extended Data Fig. 1**). Short video clips containing behavioral examples are sampled from experimental recordings and labeled with behaviors of interest, such as rearing, grooming, ect. These clips are processed by a frozen VideoPrism backbone, which extracts spatiotemporal patch token embeddings *X* ∈ R*^T^*^×*S*×*D*^, where T denotes the number of frames, S the spatial tokens per frame, and D the embedding dimensionality. Rather than fine-tuning the foundation model, tokens are grouped by frame and passed through a per-frame spatial attention pooling module that learns attention weights (α) over the S spatial tokens, producing a single frame-level embedding (h*t*) per frame. The resulting sequence of frame embeddings (h*1*, h*2*, …, h*T*) is then processed by a temporal decoder consisting of temporal pooling and a multi-stage temporal convolutional network (MS-TCN), which outputs per-frame behavioral predictions as sigmoid or softmax class scores (**Fig. 1a**). This architecture yields frame-resolved behavior sequences and bout-segmented timelines, enabling continuous segmentation of recordings into sequences of labeled actions (**Extended Data Fig.2; Supplementary Video 1**). By operating entirely on frozen embeddings, this design allows rapid customization of behavior sequencing models across experiments without retraining the underlying video model.

**Figure 1.**
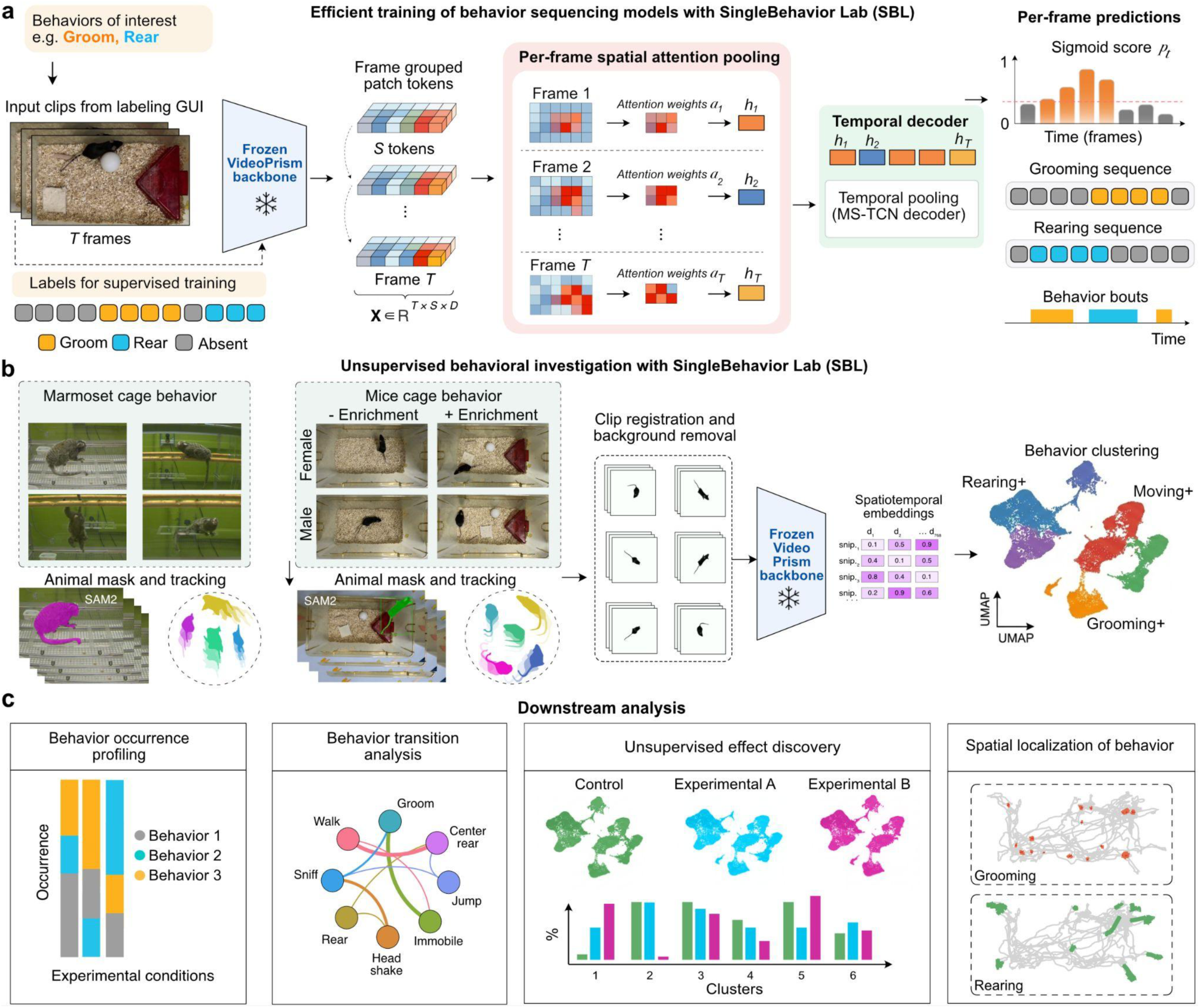
Overview of the SingleBehavior Lab (SBL) framework for efficient behavior sequencing and unsupervised behavioral discovery. a) Efficient training of behavior sequencing models. Annotated video clips (T frames) with frame level behavioral labels (e.g., Groom, Rear) are processed by a frozen VideoPrism backbone, which extracts spatiotemporal patch token embeddings. Tokens are grouped by frame and passed through a per-frame spatial attention pooling module that learns attention weights (α) over the S spatial tokens to produce a single frame level embedding per frame. The sequence of frame embeddings is decoded by a temporal decoder consisting of temporal pooling and a multi-stage temporal convolutional network (MS-TCN), yielding per-frame behavioral predictions as sigmoid or softmax class scores. The model outputs frame resolved behavior sequences and bout segmented timelines. b) Illustration of unsupervised investigation of behavior and experimental discovery. Examples of marmoset and mouse cage videos across experimental conditions are segmented and tracked using SAM2 to generate animal masks. Tracked clips undergo registration and background removal before being processed by the frozen VideoPrism backbone to extract spatiotemporal embeddings. Embeddings are clustered and visualized via UMAP, revealing emergent behavioral states (e.g., Rearing+, Moving+, Grooming+) without predefined behavioral labels. c) Illustration of downstream analysis modules. SBL supports behavior occurrence profiling across experimental conditions, behavior transition analysis via state transition graphs, unbiased effect discovery through cluster level comparisons across experimental groups (e.g., Control, Experimental A, Experimental B), and spatial localization of behavior by mapping classified actions onto tracked animal trajectories.

**Figure 2.**
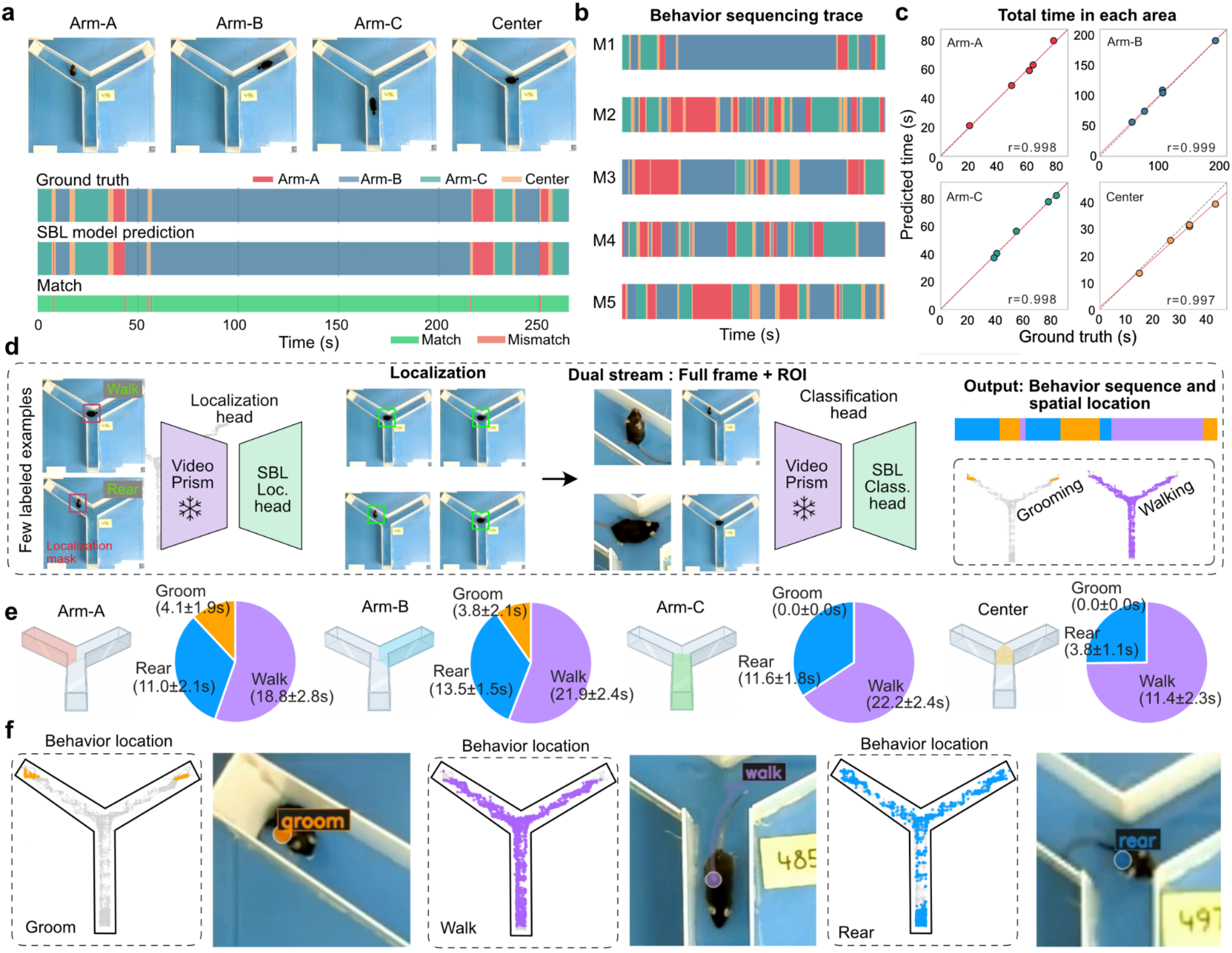
SingleBehavior Lab (SBL) enables behavior sequencing and spatial localization. **a)** SBL model prediction of behavior sequences in a three arm Y-maze (Arm-A, Arm-B, Arm-C, Center). Top, representative frames from each region. Middle, ground truth area annotations over time. Bottom, SBL predictions and match/mismatch visualization, demonstrating close agreement between predicted and annotated occupancy. **b)** Predicted behavioral sequencing traces for five mice (M1–M5), showing temporal transitions between maze regions and individual exploration strategies. **c)** Quantification of total time spent in each region. Predicted durations closely match ground truth across all areas (Pearson r = 0.997–0.999), indicating accurate temporal segmentation and area assignment. **d)** Dual-stream architecture for joint localization and classification. A pretrained VideoPrism backbone feeds a localization head to predict spatial masks and regions of interest (ROIs). Full-frame and ROI crops are processed by a classification head to generate behavior labels and temporally resolved sequences. Example outputs illustrate accurate localization of walking and rearing and integrated behavior sequencing. **e)** Region-specific behavioral quantification showing % of total duration (s) of grooming, walking, and rearing within each maze compartment, revealing spatial modulation of behavioral states. **f)** Spatial distributions of behavioral clusters across the maze. Grooming events are spatially confined, walking spans arm trajectories, and rearing localizes near junctions and boundaries, illustrating how discovered motifs map onto physical space. Error represented as SEM and n=5 mice.

To assess the efficiency of this few-shot learning strategy, we benchmarked SBL classification strategy using the Jhuang et al. mouse behavior database (MIT dataset) [33], which comprises eight annotated behavioral classes (**Extended Data Fig. 3a**). We compared three adapter training objectives on top of frozen VideoPrism embeddings: standard cross-entropy, supervised contrastive learning (SupCon), and SupCon with augmentation, across three labeling budgets of 5, 10, or 20 clips per class, corresponding to as little as 2.7 seconds of labeled video per behavior class (**Extended Data Fig. 3a**). Contrastive adapter training substantially improved few-shot classification accuracy relative to cross-entropy, with the largest gains under the most data limited conditions. With only 5 clips per class, macro F1 increased from 40.68% (baseline) to 80.76% with SupCon and augmentation (Δ = 40.1%; **Extended Data Fig. 3c–d**). Per-class analysis revealed that gains were not equal for all behavior classes: rearing improved from 14.7% to 83.5% and hanging from 36.0% to 94.0% under the same condition (**Extended Data Fig. 3b**). As the labeling budget increased to 10 or 20 clips per class, performance approached saturation and the advantage of contrastive training diminished. This indicates that contrastive objectives are most valuable when supervision is sparse, or should be used until the first version of the model is created before using that model for active learning and training data mining implemented in the SBL interface. Critically, these classification gains translated directly into improved temporal behavior sequencing, confirming that the few-shot adapter with temporal decoder are sufficient to produce temporally consistent behavior sequences suitable for downstream analysis.

In addition to supervised behavior sequencing, SingleBehavior Lab supports unsupervised exploration of behavioral structure using the same video embedding backbone (**Fig. 1b; Extended Data Fig 4**). To ensure robustness across diverse recording environments, videos are first processed using a promptable segmentation and tracking approach that extracts object centered clips of individual animals. This step enables consistent behavior analysis across different experimental setups, such as mouse home cage recordings under differential enrichment conditions and marmoset cage behavior. The extracted clips are spatially aligned and background reduced before being encoded using the frozen VideoPrism model to obtain spatiotemporal embeddings. These embeddings capture posture, motion, and temporal dynamics directly from video without requiring explicit pose estimation. Low dimensional projection of these embeddings followed by clustering reveals discrete behavioral states, representing recurrent patterns of animal activity such as locomotion, rearing, or grooming like behaviors (**Fig. 1b**). Because embeddings are derived from a shared foundation model, behavioral states can be compared across experimental contexts and species, which we demonstrate further in the paper.

SingleBehavior Lab further provides a set of downstream analysis tools that convert discovered behaviors into quantitative readouts (**Fig. 1c**). Behavior occurrence profiling quantifies the relative frequency of behavioral states across experimental conditions, enabling direct comparisons between groups. To investigate the temporal organization of actions, the framework also supports behavior transition analysis, which represents behavioral sequences as networks of state transitions using Markov Chain. These analyses capture structured patterns of behavioral progression, revealing how animals move between behavioral states over time. For experimental comparisons, we implemented unbiased effect discovery, which identifies behavioral clusters that differ between different animals, groups or experimental conditions, implementing robust metadata management systems within SBL. This approach highlights behavioral states that are selectively enriched or depleted following experimental manipulations. Furthermore, SingleBehavior Lab enables spatial localization of behaviors by mapping behavioral segments back onto animal trajectories within the arena, allowing identification of behaviors associated with specific spatial regions (**Fig. 1c; Extended Data Fig. 4**).

### Behavior sequencing and spatial localization with a unified model

To test whether SingleBehavior Lab can jointly resolve where an animal is and what it is doing, we applied the framework to a three arm Y-maze assay and trained a unified model for simultaneous area localization and behavior classification (**Fig. 2**). We used an open source video dataset obtained from Harvard Dataverse collected by L. Robinson [34]. Rather than training separate pipelines for tracking, region assignment and action recognition, we implemented a dual-stream architecture built on a shared pretrained VideoPrism backbone, enabling end-to-end optimization of spatial and behavioral outputs. We first evaluated automated area classification (Arm-A, Arm-B, Arm-C and Center) against frame-level manual annotations (**Fig. 2a**). The localization head predicts spatial masks and region-of-interest (ROI) coordinates directly from video frames, which are temporally aligned with behavioral labels. Model predictions closely matched ground-truth annotations across the full recording, with minimal mismatch epochs (**Fig. 2a**). Temporal sequencing traces across five animals (**Fig. 2b**) revealed coherent region transitions and individual exploration strategies, demonstrating stable frame-to-frame predictions. Quantitative comparison of predicted versus annotated occupancy times showed high agreement across all maze regions (Pearson r = 0.997–0.999; **Fig. 2c**), confirming that the model accurately segments spatial states over time. Importantly, this performance was achieved without task specific backbone fine-tuning, relying instead on lightweight trainable heads.

The computational architecture integrates localization and classification through a dual-stream design (**Fig. 2d**). A shared VideoPrism encoder extracts spatiotemporal features that are routed to a localization head producing spatial masks and ROIs and a classification head operating on both full-frame embeddings and cropped ROIs. This design allows the classifier to leverage global contextual cues (maze geometry) together with fine grained posture information from the ROI. Since the backbone is pretrained on 288×288 pixel inputs, the same resolution used by SBL during embedding extraction, full-frame processing alone can lead to loss of visual detail needed for resolving subtle behaviors. The dual-stream design addresses this by providing the classification head with both wide context and fine grained ROI information, ensuring that spatial attention and behavioral discrimination are mutually informative. To verify that the localization head converges reliably under few-shot supervision, we registered training loss, center error, and Intersection over Union (IoU) across epochs (**Extended Data Fig. 5a–c**). Training loss and center error decreased rapidly within the first five epochs and plateaued at low values, while IoU stabilized above 0.90, confirming that accurate animal localization is achieved with minimal labeled masks (**Extended Data Fig. 5b-c**). Qualitative inspection of predicted crops across epochs further confirmed that precise centering on the animal was reached by approximately epoch 10 and maintained thereafter (**Extended Data Fig. 5d**). Together, these results demonstrate that the localization head learns spatially precise animal crops with few-shot supervision, enabling reliable ROI extraction for downstream classification. To verify convergence of the classification head, we tracked training accuracy, validation loss, and per-class F1 scores across epochs (**Extended Data Fig. 6a–d**). Both accuracy and validation loss stabilized rapidly, with all three behavioral classes (rear, groom, walk) reaching F1 scores above 95% by epoch 6, confirming that few-shot supervision is sufficient for robust behavioral discrimination (**Extended Data Fig. 6c, d**). We next quantified region specific behavioral structure (**Fig. 2e**). This analysis revealed consistent and compartment specific distributions of grooming, walking, and rearing across the Y-maze. Walking dominated all compartments, but dwell times differed markedly across zones. Animals spent 18.8 (± 2.8)s walking in Arm-A, 21.9 (± 2.4)s in Arm-B, and 22.2 (± 2.4)s in Arm-C, compared to only 11.4 (± 2.3)s in the Center, consistent with reduced locomotion in the spatially constrained central junction. Rearing was similarly distributed across arms (Arm-A: 11.0 ± 2.1s; Arm-B: 13.5 ± 1.5s; Arm-C: 11.6 ± 1.8s) but was substantially reduced in the Center (3.8 ± 1.1s), suggesting that exploratory vertical behavior is preferentially expressed in the arm compartments. Grooming showed the most spatially restricted pattern, it was detected exclusively in Arm-A (4.1 ± 1.9s) and Arm-B (3.8 ± 2.1s), with no grooming events recorded in Arm-C or the Center (0.0 ± 0.0s for both), indicating a strong spatial bias of self directed behavior toward specific arms. Behavior sequencing traces show distribution of each behavior class over time (**Extended Data Fig. 6e; Supplementary Video 2**). Together, these results demonstrate that integrated localization and behavioral sequencing outputs support region specific quantification of behavioral states.

Finally, mapping clustered behavioral events back onto spatial trajectories demonstrated interpretable spatial organization (**Fig. 2f**). Grooming events were spatially restricted, walking spanned entire arm trajectories, and rearing localized near arm junctions and boundaries. Critically, this spatial mapping is enabled by the localization head, which predicts frame-by-frame animal centroid coordinates directly from the shared backbone features. These coordinates provide continuous tracking of the animal’s position over time without an external tracking module. It is important to mention that this is a different approach fully integrated into the SBL model, in contrast to using SAM2 for tracking (also implemented here). Because localization and behavior classification are trained within the same model, each predicted behavioral label is inherently aligned to precise spatial coordinates. This enables trajectory level reconstruction of specific behaviors, allowing behavioral states to be projected onto the animal’s movement path across the maze. To further assess what spatial features the model uses for behavioral discrimination, we visualized attention maps during classification of individual behaviors (**Extended Data Fig. 7**). For grooming, attention concentrated on the animal’s head and forepaws within the ROI; for walking, attention was distributed along the body axis and limbs; and for rearing, the model attended to the upright body posture and the animal’s position relative to the maze walls (**Extended Data Fig. 7a–c**). Projecting these attention maps back onto full frame inputs confirmed that the model selectively focuses on behaviorally relevant body regions while suppressing irrelevant background features. These results confirm that the SBL models learn interpretable, behavior specific spatial representations rather than relying on contextual shortcuts. Together, this unified architecture of SBL couples frame resolved tracking with action sequencing, supporting integrated spatiotemporal analysis within a single computational framework.

### Motion aware tracking and few-shot behavior sequencing in individual drosophila

To assess whether SingleBehavior Lab can be applied to smaller animals and denser multi animal recordings, we next analyzed drosophila behavior in a circular arena containing multiple animals recorded simultaneously from openly available dataset [35] (**Fig. 3**). First, we integrated a motion aware optimization layer on top of SAM2 within the SBL framework, enabling robust segmentation and identity preserving tracking of multiple flies simultaneously in a circular open field arena. The approach successfully resolved all 12 individuals across the recording, producing segmentation masks and reconstructed spatial trajectories that captured the full extent of each animal’s movement within the arena (**Fig. 3a**). Trajectories revealed substantial inter individual variability in spatial exploration patterns, with some flies exhibiting strong wall-following behavior while others crossed interior regions, consistent with known individual differences in thigmotaxis and locomotor activity in drosophila.

**Figure 3.**
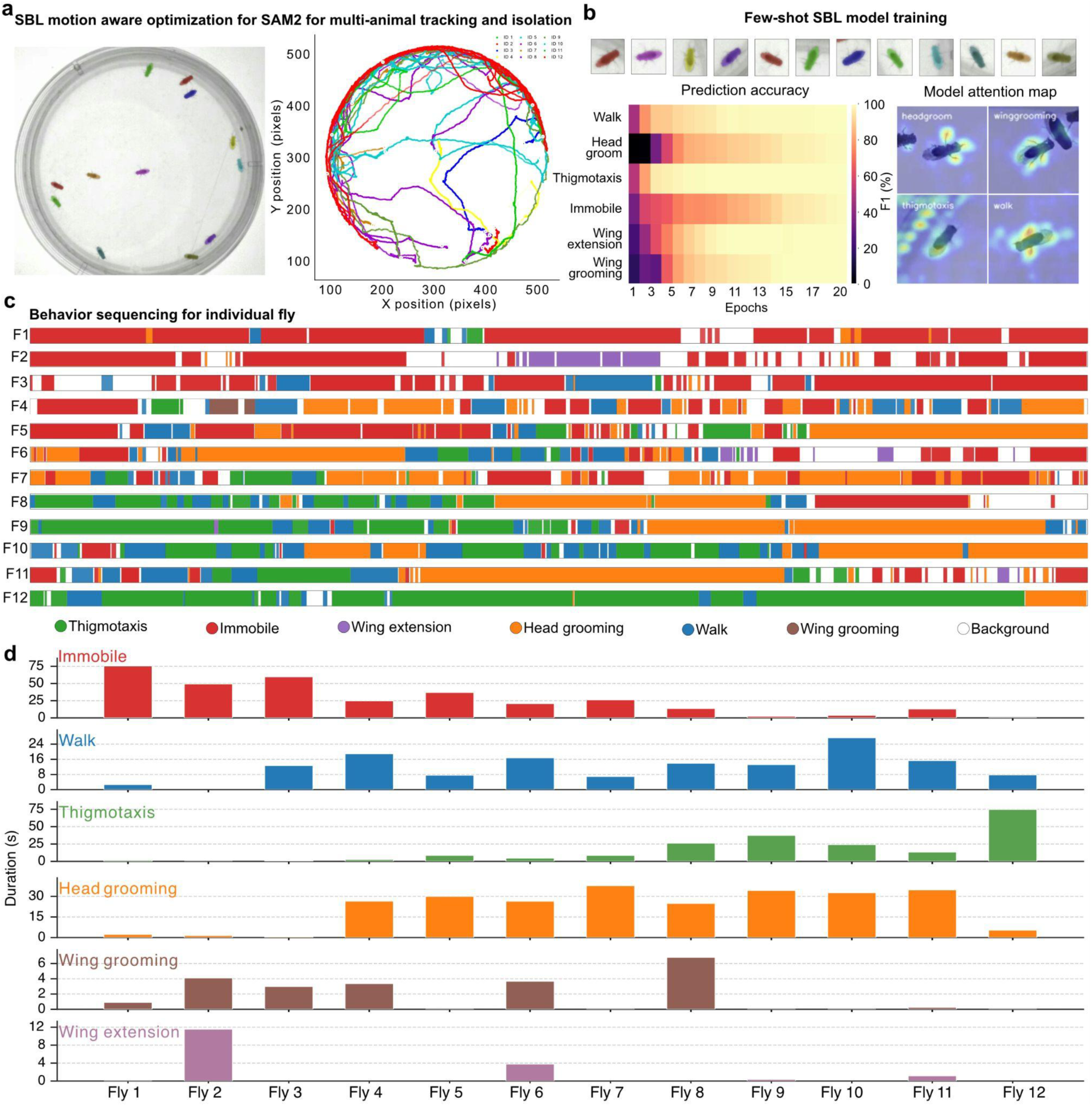
Motion aware segmentation and behavior sequencing in individual flies. **a)** Motion-aware optimization built on SAM2 within SingleBehavior Lab (SBL) for isolating and tracking multiple individual flies in a circular arena. Left: representative frame showing segmented and color labeled flies. Right: reconstructed trajectories for individually tracked flies plotted in arena coordinates. **b)** Few-shot behavior classification using SBL. Top: representative fly crops used as model input obtained from tracking centroids. Bottom left: heatmap showing prediction accuracy on unseen data across training epochs for individual behavior classes. Bottom right: representative model attention maps for selected classes. **c)** Predicted behavior sequences for individual flies (F1–F12) shown as temporally ordered behavioral timelines, with colored segments denoting class labels across each recording. **d)** Quantification of total behavior duration per individual fly across the full recording, displayed as separate panels for each of the six behavior classes. Each panel shows cumulative duration in seconds on the y-axis for individual flies (F1–F12) on the x-axis. F: fly.

To specifically get insights into behavioral states from tracked individuals, we asked whether SBL could achieve accurate action recognition from minimal labeled examples. Using centroid aligned fly crops extracted directly from the tracking output, we trained a lightweight SBL model on top of frozen VideoPrism embeddings under a few-shot regime. Prediction accuracy, assessed by per-class F1 score across training epochs, increased rapidly for most behavioral classes, including walking, immobility, and thigmotaxis, reaching high performance within the first few epochs (**Fig. 3b, bottom left**). Rarer or more morphologically subtle behaviors, including head grooming and wing grooming, required additional epochs but ultimately achieved reliable prediction. Gradient weighted attention maps confirmed that the model focused on behaviorally relevant spatiotemporal features, including body posture and appendage configuration, while for thigmotaxis the model gives partial attention to edges of the dish (**Fig. 3b, bottom right**), supporting the interpretability of the learned representations.

Applying the trained classifier to full recording sessions produced temporally resolved behavioral sequences for each of the 12 tracked flies (**Fig. 3c; Supplementary Video 3**). Individual behavioral timelines revealed marked inter-individual heterogeneity in both the identity and sequential structure of expressed behaviors. Some flies spent the majority of the session engaged in thigmotaxis or locomotion (e.g., F9, F12), while others exhibited more complex sequences with frequent transitions between grooming, immobility, and walking (e.g., F6, F7). F1 appeared predominantly immobile throughout the session, illustrating SBL’s sensitivity to outlier behavioral phenotypes that would be obscured in group-level analyses. To quantify this variability, we computed the cumulative duration of each behavior class per individual across the full recording (**Fig. 3d**). This analysis confirmed substantial fly-to-fly differences in time allocation across all six categories. Together, these results demonstrate that SBL’s motion aware segmentation and few-shot behavior sequencing pipeline supports individual level resolution behavioral phenotyping in multi-animal recording contexts.

### Unsupervised behavioral clustering reveals behavioral phenotypes in home cage recordings

To test whether SingleBehavior Lab can discover behavioral structure without predefined labels, we applied the unsupervised analysis pipeline to home-cage recordings of male and female mice housed under standard or enriched conditions (**Fig. 4a**). Animals were segmented and tracked using point-prompt SAM2 through the SBL GUI, and single-animal clips were registered and background-removed before embedding extraction (**Fig. 4b**). Spatiotemporal embeddings were computed using the frozen VideoPrism backbone and clustered using the Leiden algorithm within the SBL clustering interface, which provides configurable controls for UMAP dimensionality reduction, clustering algorithms and resolution, and direct video clip preview for each data point (**Extended Data Fig. 4a**). Low-dimensional visualization via UMAP revealed well-separated behavioral clusters, which we annotated by reviewing automatically extracted representative video clips from each cluster using the SBL interface (**Fig. 4c**). Cluster names carry a “+” suffix to indicate that each cluster is enriched for a given behavior rather than exclusively containing it, reflecting the continuous and overlapping nature of naturalistic actions.

**Figure 4.**
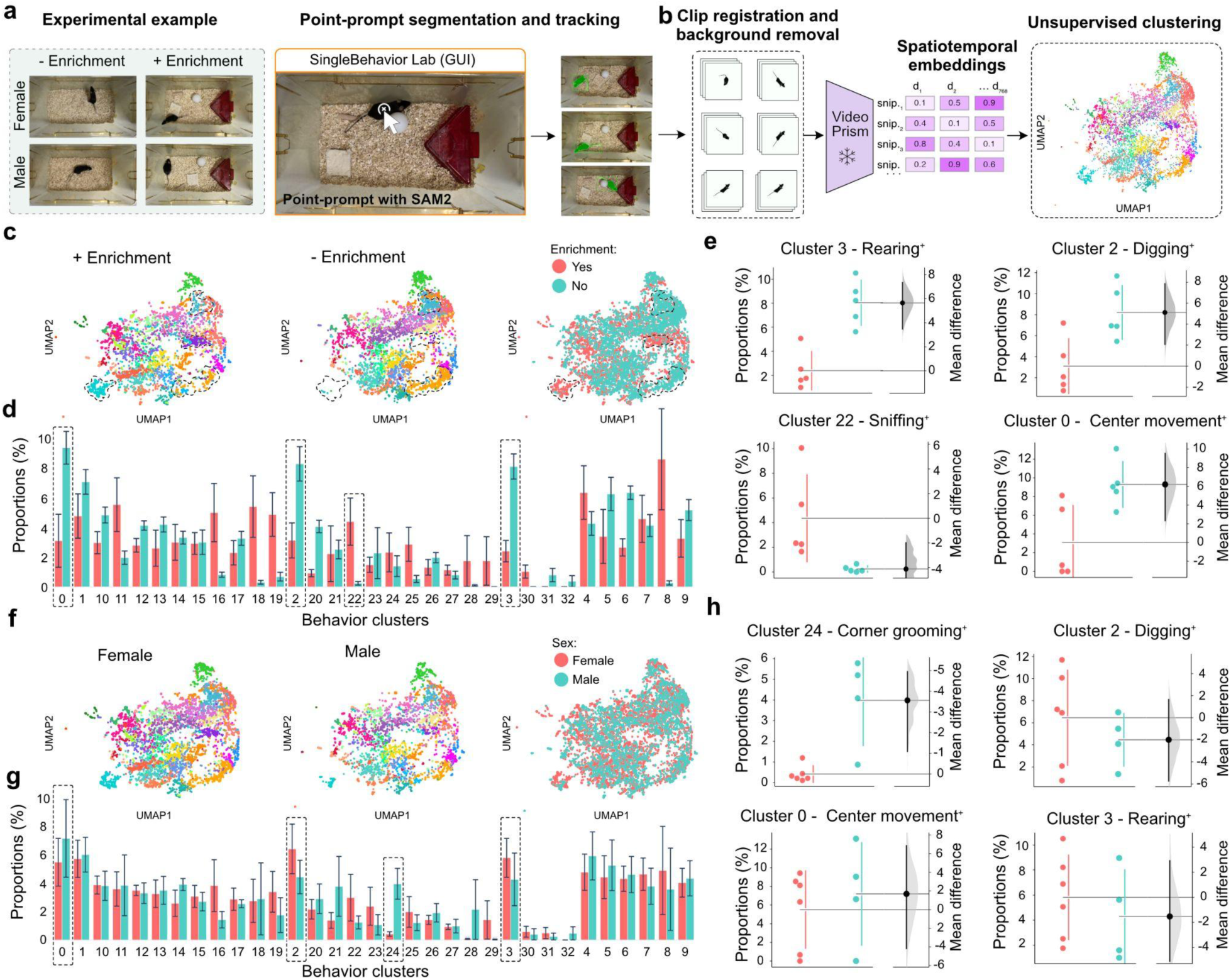
SingleBehavior Lab enables unsupervised behavioral clustering and discovery in mouse home-cage recordings. **a)** Experimental setup and animal tracking pipeline. Home-cage recordings from male and female mice under enrichment and -Enrichment conditions were processed using point-prompt segmentation and tracking with SAM2, implemented through the SingleBehavior Lab GUI. **b)** Unsupervised analysis workflow. Registered and background-removed single-animal clips were passed through the frozen VideoPrism backbone to extract spatiotemporal embeddings. Embeddings were clustered and visualized using UMAP, revealing distinct behavioral states without predefined behavioral labels. Cluster names reflect the predominant behavior enriched within each cluster; the “+” suffix denotes that clusters are not behaviorally pure and represent enrichment of a given behavior rather than its exclusive occurrence. **c)** UMAP embeddings of single-behavior units (SBUs) under enrichment and -Enrichment conditions (including males and females), colored by Leiden cluster identity. Named clusters (e.g., Rearing, Digging, Sniffing, Center movement) were identified by reviewing automatically extracted representative video clips from each cluster using the SBL graphical interface. **d)** Behavior cluster quantification by enrichment condition. Bar plots show the mean fraction of SBUs assigned to each behavior cluster per animal, compared between enrichment and -Enrichment groups. Dashed boxes indicate clusters with notable differences between conditions. **e)** Estimation plots for representative behavior clusters comparing enrichment conditions (Rearing+, Digging+, Sniffing+, Center movement+). Each dot represents one animal. Violin plots show the bootstrapped distribution of mean differences, with the point estimate and 95% confidence interval indicated. **f)** UMAP embeddings of single-behavior units (SBUs) by sex (female: pink vs male: teal; with and without enrichment condition), showing the joint distribution of behavioral phenotypes across the two grouping variables. **g)** Behavior cluster quantification by sex. Bar plots show the mean fraction of SBUs per behavior cluster compared between female and male animals across conditions. Data are shown as mean ± s.e.m. Dashed boxes indicate clusters with notable sex differences. **h)** Estimation plots for representative behavior clusters comparing sexes (Corner grooming+, Digging+, Center movement+, Rearing+). Each dot represents one animal. Data are shown as mean ± s.e.m. Violin plots show the bootstrapped distribution of mean differences between groups, with the point estimate and 95% confidence interval indicated.

Clustering of the joint embedding space identified a diverse repertoire of behavioral states, including rearing, digging, sniffing, center-cage movement, and corner grooming, among others (**Fig. 4c**). The relative abundance of each cluster across experimental groups could be directly compared through the cluster proportions view (**Extended Data Fig. 4b**), while the spatial cluster distribution view enabled mapping of behavioral clusters onto the animal’s position within the arena (**Extended Data Fig. 4c**). To assess whether environmental enrichment altered behavioral composition, we separated the joint embedding by condition and quantified the fraction of single-behavior units (SBUs) assigned to each cluster per animal (**Fig. 4c–d**). Several clusters showed visible enrichment-dependent shifts in occupancy. To quantify these effects, we examined representative clusters using estimation statistics (**Fig. 4e**). Rearing+ (Cluster 3) and Digging+ (Cluster 2) showed altered occupancy between enrichment and without enrichment groups, as did Sniffing+ (Cluster 22) and Center movement+ (Cluster 0). These results indicate that environmental enrichment systematically reshapes the behavioral repertoire, with specific motor and exploratory behaviors differentially affected.

We next asked whether sex contributed to variation in behavioral phenotype independently of enrichment conditions. Projecting the joint embedding separately for female and male animals (pooled across enrichment conditions) revealed broadly overlapping behavioral landscapes with localized sex-dependent differences (**Fig. 4f**). Quantification of cluster occupancy by sex (**Fig. 4g**) identified several clusters with divergent representation between males and females. Estimation plots for representative clusters (**Fig. 4h**) showed that Corner grooming+ (Cluster 24) and Digging+ (Cluster 2) differed between sexes, as did Center movement+ (Cluster 0) and Rearing+ (Cluster 3). These sex-linked behavioral differences complement the enrichment effects and suggest that the unsupervised embedding captures biologically meaningful variation along multiple experimental axes.

Together, these results demonstrate that SingleBehavior Lab can extract interpretable behavioral states from naturalistic home-cage recordings in a fully unsupervised manner. The combination of foundation model embeddings and graph-based clustering enables detection of condition and sex-dependent behavioral phenotypes without requiring manual behavior definitions, offering a scalable approach for unbiased behavioral screening in experimental neuroscience.

### Multi-animal tracking enables scalable unsupervised behavioral discovery

Having established unsupervised behavioral discovery in single-animal recordings (**Fig. 5**), we next asked whether SingleBehavior Lab generalizes to multi-animal settings while preserving individual identities and enabling joint embedding analysis. We applied the framework to home-cage recordings containing two freely interacting animals per arena (**Fig. 5a**). Using point-prompted SAM2 within the GUI, each animal was segmented and tracked across frames, generating identity-preserved masks and trajectories without manual keypoint annotation. This enabled extraction of behavior snippets independently for each animal despite shared space and partial occlusions.

**Figure 5:**
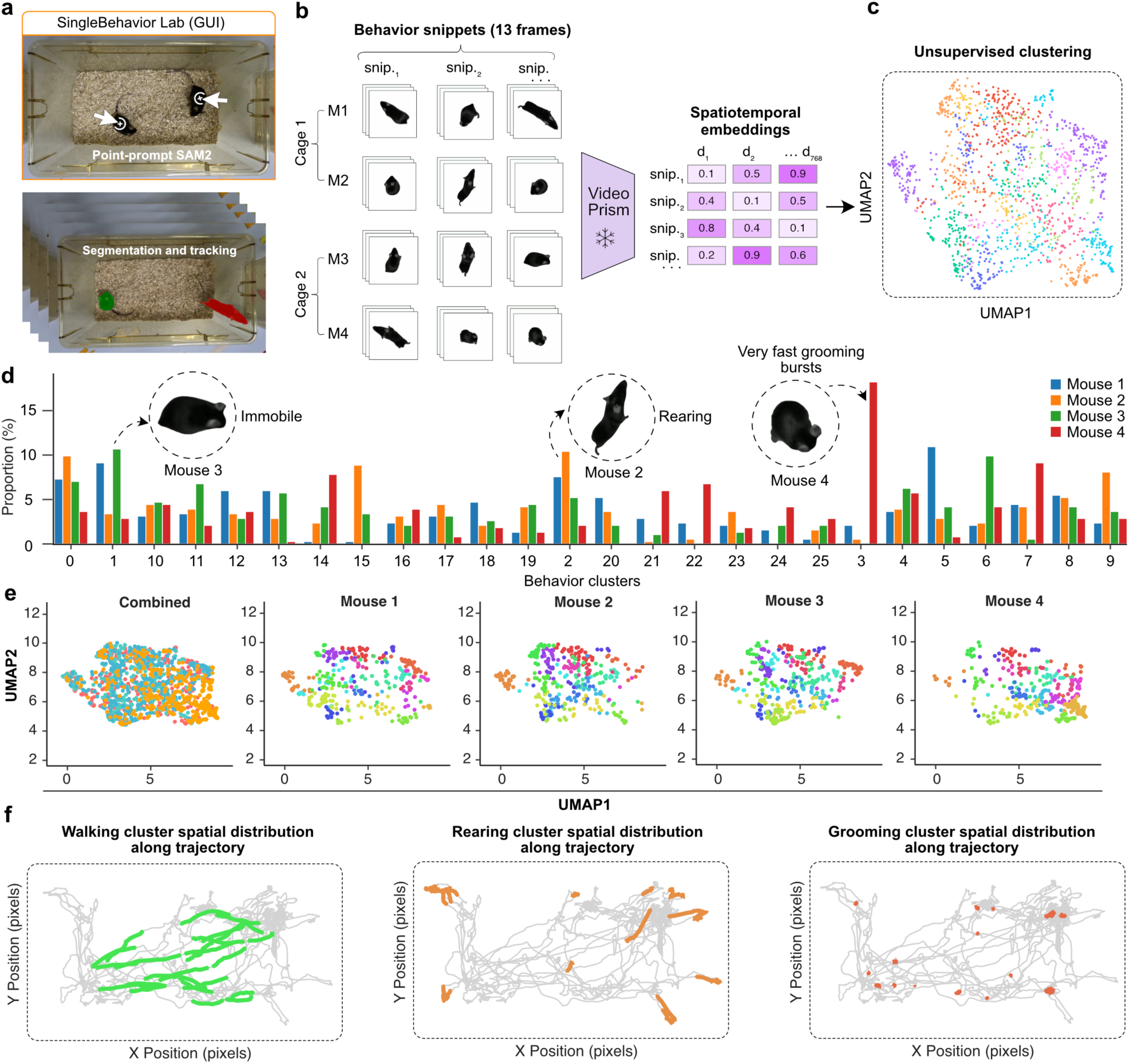
Multi-animal behavior embedding integration and analysis. **a)** Graphical user interface (GUI) of SingleBehavior Lab. Top: point-prompted SAM2 enables robust object segmentation in multi-animal arenas. Bottom: automated segmentation and tracking generate identity-preserved trajectories and cropped behavioral clips across contexts. **b)** Behavioral snippet extraction and embedding. Short video snippets (13 frames) are sampled from tracked individuals across cages and mice (M1–M4). Snippets are passed through a pretrained foundation video model (VideoPrism) to generate high-dimensional spatiotemporal embeddings (d₁–d₇₆₈). These embeddings form the basis for downstream clustering and adapter training. **c)** Unsupervised clustering of snippet embeddings visualized with UMAP. Each point represents a behavioral snippet, revealing structured behavioral motifs without manual labels. **d)** Proportion of time spent in discovered clusters across mice. Cluster identities correspond to interpretable behaviors (examples shown: immobility, rearing, very fast grooming bursts). Bar plots illustrate inter-animal differences in behavioral composition. **e)** UMAP visualization of embeddings colored by cluster identity for all mice combined and for each individual mouse, demonstrating shared and individual-specific behavioral structure. **f)** Spatial distribution of selected behavioral clusters along animal trajectories. Walking clusters distribute broadly across the arena, rearing events localize to specific regions, and grooming bursts appear as spatially sparse events, illustrating how discovered motifs map onto physical space.

Continuous tracks were partitioned into short spatiotemporal snippets (13 frames), which were embedded using the pretrained VideoPrism backbone to generate high-dimensional representations (**Fig. 5b**). These embeddings were pooled across cages and individuals and subjected to UMAP projection and Leiden clustering without behavioral supervision. The resulting embedding space exhibited well-separated and compact clusters corresponding to discrete behavioral motifs (**Fig. 5c**), demonstrating that pretrained video features retain discriminative structure even in multi-animal contexts.

Cluster occupancy profiles revealed both shared and individual-specific behavioral structure (**Fig. 5d**). While core motifs such as immobility and rearing were consistently detected across animals, their relative proportions differed between individuals, including rare high-frequency grooming bursts enriched in specific mice. Visualization of the joint embedding alongside per-animal projections showed that clusters remained geometrically coherent across subjects (**Fig. 5e**), indicating that the learned representation captures common behavioral primitives while preserving inter-animal variability.

Because tracking is integrated with segmentation in the same pipeline, each embedded snippet retains precise spatial coordinates for the corresponding individual. Mapping cluster identities back onto trajectories revealed interpretable spatial organization of behaviors (**Fig. 5f**). Walking motifs spanned extended arena paths, rearing events localized near boundaries, and grooming clusters appeared as spatially confined events. Importantly, these trajectory-level reconstructions are derived directly from identity-preserved tracking outputs rather than post-hoc alignment procedures.

Together, these results demonstrate that SingleBehavior Lab scales from single- to multi-animal recordings, enabling identity-resolved tracking and unsupervised behavioral discovery within a unified computational framework. The combination of multi-animal segmentation, foundation-model embeddings and clustering supports unbiased extraction of shared and individual-specific behavioral motifs in socially relevant contexts.

### SingleBehavior Lab enables integrated behavior sequencing and unsupervised behavioral discovery

To demonstrate the generalizability of SingleBehavior Lab across species and recording conditions, we applied the framework to continuous cage recordings of common marmosets (**Data collection: Methods [36]**). We first assessed whether SBL could produce temporally resolved behavioral sequences from extended marmoset recordings. A 20-minute continuous session was divided into four successive 5-minute video segments, and the SBL behavior sequencing model was trained to classify seven behavioral states: Hang, Scratch, Jump, Groom, Rear, Walk, and Stationary (**Fig. 6a**).

**Figure 6.**
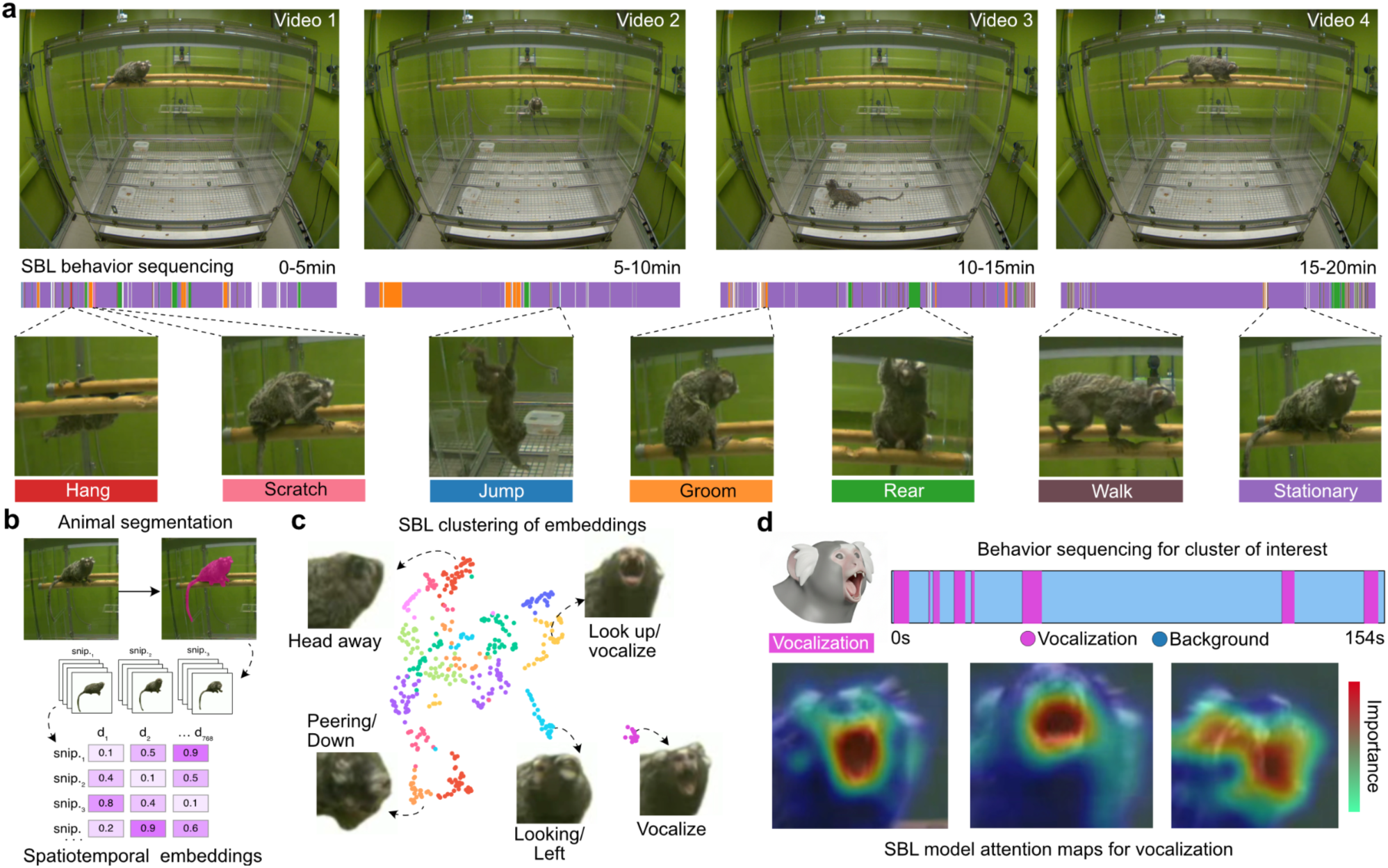
SingleBehavior Lab integrates automated behavior sequencing with unsupervised spatiotemporal embedding clustering exemplified in marmoset cage behavior. **a)** SBL behavior sequencing applied to a continuous 20-minute marmoset cage recording, divided into four successive 5-minute video segments. Color-coded behavioral timelines indicate the temporal distribution of annotated behavioral states, including Hang, Scratch, Jump, Groom, Rear, Walk, and Stationary, across the full recording session, revealing dynamic shifts in behavioral composition over time. **b)** Schematic of the SBL object segmentation and embedding pipeline. Animals are segmented from background using SAM2-based video object segmentation, yielding object-centered clips (snips). Each snip is passed through the frozen VideoPrism backbone to extract high-dimensional spatiotemporal embedding vectors (d₁–d₃₆₈), forming the basis for downstream unsupervised analysis. **c)** Unsupervised clustering of spatiotemporal embeddings reveals discrete behavioral motifs in marmoset cage behavior. Embeddings from individual snips are projected into a low-dimensional space and grouped into clusters corresponding to interpretable behavioral states, including Head away, Peering/Down, Looking/Left, Vocalize, and Look up/Vocalize. Representative frames illustrate the visual content of each cluster. White frames represent background frames that are ignored by the model. **d)** Cluster-resolved behavior sequencing for a behavioral state of interest (Vocalization). The temporal sequence of vocalization events is visualized across a 154-second window, highlighting the episodic structure of this behavior. SBL model attention maps (bottom) display the spatiotemporal regions driving model predictions.

Beyond whole-body behavior sequencing, we asked whether SBL could resolve finer-grained, head-related behavioral states by directing the segmentation pipeline to focus on a specific body region. Using the SAM2 point-prompt segmentation within SBL, we targeted the marmoset’s face rather than using full frame video, generating face-centered clips from the cage recordings (**Fig. 6b**). Face-centered video was processed through the frozen VideoPrism backbone to extract spatiotemporal embedding vectors capturing facial posture, gaze direction, and mouth dynamics (**Fig. 6b**). Projection of these embeddings into a low-dimensional space followed by clustering revealed discrete head-related behavioral states that would be difficult to resolve from full-body video alone, including Head away, Peering/Down, Looking/Left, Vocalize, and Look up/Vocalize (**Fig. 6c**). Notably, two distinct vocalization-related clusters emerged (Vocalize and Look up/Vocalize) showing that the spatiotemporal embeddings encoded not only the vocal behavior itself but also the accompanying head posture. These results demonstrate that the flexible segmentation strategy in SBL allows users to guide the level of behavioral resolution by choosing where to focus, enabling discovery of subtle behavioral states that depend on specific anatomical regions.

To explore how discovered facial behavioral states unfold over time, we applied the SBL behavior sequencing approach. Focusing on vocalization events, we visualized the temporal sequence across a 154-second window, revealing the episodic structure of vocal behavior interspersed with background periods (**Fig. 6d**). To examine what spatial features drove the model’s predictions, we inspected the SBL attention maps for vocalization (**Fig. 6d**). The attention maps concentrated on the marmoset’s face and open mouth, the anatomically relevant regions for vocal production, confirming that the model learned biologically meaningful feature representations. This interpretability was consistent across other behavioral classes: attention maps for grooming concentrated on the head and forepaws, walking attention distributed along the body axis and limbs, rearing attention captured upright body posture, and turning attention focused on the rotational axis of the body (**6d**). Together, these results demonstrate that SBL generalizes across species and that its segmentation-guided embedding strategy and behavior sequencing can be combined to enable discovery of both coarse whole-body behaviors and fine-grained head and face-related behavioral states, with interpretable attention maps confirming that predictions are grounded in anatomically relevant features.

## Discussion

Here we present SingleBehavior Lab (SBL), a unified framework for behavioral analysis that combines few-shot behavior sequencing with unsupervised behavioral discovery, built on frozen video foundation model embeddings. By decoupling visual feature extraction from task-specific behavioral decoding, SBL enables rapid adaptation to new species, recording conditions, and behavioral vocabularies without retraining the underlying video model. We demonstrated the framework across flies, mice and marmosets, in Y-maze, home-cage, and enrichment paradigms, showing that a single architectural design supports frame-resolved behavior sequencing, joint spatial-behavioral analysis, unsupervised behavioral clustering, and region-specific behavioral discovery through flexible segmentation. A central design principle of SBL is that behavioral classification should be achievable with minimal labeled data. Our benchmarking on the Jhuang et al. mouse behavior database [33] showed that contrastive adapter training on frozen VideoPrism embeddings reached 80.76% macro F1 with as few as 5 labeled clips per class, corresponding to under 3 seconds of annotated video per behavior. This performance was sufficient to produce temporally consistent behavior sequences with bout boundaries closely matching human annotations. In practice, we found that the quality of the labeled examples matters more than their quantity. As few as 5 to 10 well-chosen, cleanly segmented 8-frame clips per behavior were often sufficient for strong classification performance, provided that the clips clearly captured the target behavior without ambiguity. Conversely, increasing the number of training examples with noisy or ambiguous labels yielded diminishing returns, suggesting that curation of a small number of high-quality examples is a more effective strategy than collecting large volumes of weakly labeled data.

The model’s classification accuracy was closely linked to the degree of behavioral ambiguity inherent in the ethogram. Behaviors that are well-defined and visually distinct, such as hanging or walking, were classified with high accuracy even under extreme data limitation. In contrast, behaviors that are inherently ambiguous or graded, such as transitions between resting and grooming, or subtle postural differences between rearing and sniffing, were more difficult for the model to resolve. Importantly, this pattern mirrored the difficulty experienced by human annotators, the behaviors that were hardest for humans to agree on during manual annotation were also the behaviors most frequently confused by the model. This correspondence suggests that model errors in these cases reflect genuine ambiguity in the behavioral signal rather than a failure of the representation, and that classification boundaries in such cases are constrained by the consistency of the supervision itself. Active learning and hard-negative review within SBL provide a practical path to iteratively improve annotation quality for these ambiguous cases.

The dual-stream architecture for joint behavior classification and spatial localization demonstrated that SBL can integrate where an animal is with what it is doing within a single model. In the Y-maze, the localization head achieved near-perfect agreement with manual annotations (Pearson r > 0.99) while providing frame-aligned spatial coordinates for downstream behavioral mapping. This integration eliminated the need for separate tracking pipelines and enabled direct computation of region-specific behavioral profiles and trajectory-level behavioral reconstructions. The attention maps confirmed that the classification head learns behavior-specific spatial representations, attending to the head and forepaws during grooming, the body axis during walking, and upright posture during rearing, rather than relying on contextual or positional shortcuts. These interpretable representations increase confidence that the model’s predictions reflect genuine behavioral features.

The unsupervised analysis pipeline in SBL addresses a complementary need in behavioral neuroscience, the ability to discover behavioral phenotypes without imposing predefined categories. By clustering spatiotemporal embeddings from the same frozen backbone used for supervised analysis, SBL revealed condition-dependent and sex-dependent behavioral phenotypes in mouse home-cage recordings and species-specific behavioral motifs in marmoset cage recordings. An important consideration when interpreting these results is that behavioral clusters are not pure, each cluster is enriched for a given behavior but may contain related or transitional actions. This is an inherent property of continuous naturalistic behavior, where actions grade into one another without sharp categorical boundaries. We therefore recommend that group differences identified through unsupervised clustering be treated as hypotheses to be confirmed through targeted follow-up. Specifically, clusters of interest can be used to guide the training of dedicated behavior sequencing models, which provide frame-resolved temporal structure and quantitative bout statistics that offer deeper insight into the nature of the behavioral difference. This two-stage workflow, unsupervised discovery followed by supervised confirmation, leverages the complementary strengths of both analysis modes within SBL.

A related challenge is that some clusters resist straightforward behavioral interpretation. At high clustering resolution, the Leiden algorithm may separate behavioral states into granular subgroups that do not correspond to easily nameable actions, particularly when the underlying behavior involves continuous postural variation or context-dependent modulation. Rather than treating this as a limitation, we suggest that the appropriate clustering granularity should be determined empirically for each dataset by systematically varying the resolution parameter and evaluating cluster interpretability through the SBL video preview interface. Coarser resolutions yield broader, more easily interpretable categories, while finer resolutions can reveal subtle behavioral substructure that may carry biological significance, as demonstrated by the separation of two distinct vocalization-related clusters in marmoset face-centered embeddings.

The flexibility of the segmentation pipeline proved to be a key advantage for accessing different levels of behavioral resolution. By directing SAM2 point-prompt segmentation to the marmoset face rather than the full body, we were able to discover fine-grained head-related behavioral states, including gaze direction and vocalization posture, that would be unresolvable from whole-body video. This demonstrates that the choice of segmentation target effectively defines the scope of behavioral analysis, allowing users to move between whole-body ethograms and region-specific behavioral phenotyping within the same framework. To support robust tracking across extended recordings, we optimized the memory mechanism of the SAM2 model to enable longer and more stable segmentation without identity loss or mask degradation over time, which was critical for processing continuous multi-minute sessions typical of home-cage and cage behavior paradigms.

Several limitations and directions for future work merit consideration. First, while the frozen VideoPrism backbone provides strong general purpose representations, domain-specific fine-tuning or adaptation of the backbone on large-scale behavioral video datasets could further improve sensitivity to subtle behavioral features, particularly in species or recording environments that differ substantially from the backbone’s pretraining distribution. Second, the current active learning loop relies on user review through the GUI, future iterations could incorporate model confidence calibration to reduce the volume of clips requiring manual inspection. Finally, while we demonstrated cross-species generalizability across mice and marmosets, systematic evaluation across a broader range of species and behavioral paradigms will be important for establishing the boundaries of the framework’s applicability.

In summary, SingleBehavior Lab provides a unified, accessible platform for both hypothesis-driven behavior sequencing and hypothesis-free behavioral discovery. By combining a frozen video foundation model with lightweight trainable modules, flexible segmentation, and integrated downstream analysis tools, SBL lowers the barrier to standardized behavioral quantification across experimental contexts while maintaining the interpretability and rigor required for mechanistic neuroscience.

## Methods

### Animals

Newly-generated datasets using animals were conducted in accordance with the Regierung von Oberbayern, Germany. The mice used for the experiments were adult animals with a C57BL/6J background. Mice were between 8-16 weeks old and weighed 22-30 grams. Mice were housed in groups of up to five in enriched Type-II cage on an IVC rack system under a 12-hour day-night light cycle with ad-libitum food and water.

### Behavioral testing and used open datasets

Home cage recordings were made using a GoPro HERO12 black camera (GoPro Inc., USA) at 30 frames per second. The GoPro was mounted on a stable stand right above the cage with a distance of 36,5 cm to the bottom of the cage to provide an overview of the full cage in the frame. Male or female mice were recorded with or without enrichment. Original videos were acquired in 5K resolution and converted to 1080 HD using HandBreak (v1.10.2). To evaluate the framework across diverse behavioral paradigms and species, we used four publicly available or collaborator provided video datasets. To assess the efficiency of the few-shot learning strategy, we benchmarked SBL on the Jhuang et al. (2010) mouse behavior database [33]. Y-maze exploration recordings in rodents were obtained from L. Robison (2025) via Harvard Dataverse [34]. Multi-animal fly behavioral recordings were obtained from the UDMT dataset Li, et al. (2024) [35]. Marmoset social behavior videos were kindly provided by Dr. Kaneko and colleagues, whose dataset and experimental paradigm are described in Kaneko et al. (2024) [36].

### SBL behavior sequencing model architecture

We developed an end-to-end behavior sequencing framework for automated behavioral classification in animal video recordings. The architecture combined a frozen pretrained video backbone for spatiotemporal feature extraction with a lightweight, trainable behavior decoding head and temporal decoder. The system was designed to operate in low annotation settings with an active learning approach that enables efficient data mining and fast model optimization, resulting in framewise prediction, boundary aware sequencing, calibrated outputs, and iterative human-in-the-loop retraining.

#### Frozen video backbone

Spatiotemporal features were extracted using a frozen VideoPrism backbone. We used videoprism_public_v1_base (VideoPrism-B), which is a ViT-B backbone video encoder with 114M parameters. For each input clip of T frames, the backbone produced a tensor of patch level token embeddings of shape B x (T x S) x D, where B denoted the batch size, S the number of spatial tokens per frame, and D the embedding dimension. The backbone remained frozen throughout training, preserving the general visual representations learned during large scale pretraining and substantially reducing the number of trainable parameters. To avoid redundant computation across repeated experiments, backbone embeddings were optionally precomputed and cached to disk.

#### Data representation and supervision

Training was performed on short fixed length clips of T frames sampled from longer video recordings. Each clip could carry a combination of supervision signals: a primary clip level label, optional multi label behavior annotations, optional framewise labels, optional bounding box targets, and optional class specific hard negative metadata. This flexible annotation scheme allowed the framework to operate under both weakly supervised (clip-level only) and strongly supervised (frame-level) conditions. The system supported two output modes. In mutually exclusive multi-class mode, each frame was assigned to exactly one behavior class using a softmax output over all classes. In one-vs-rest (OvR) mode, each class was predicted independently through a sigmoid activation, which was appropriate when behaviors could co-occur or when one behavior constituted a subtype of another. Rejected predictions from model review could be stored as class-specific hard negatives, providing targeted negative supervision to suppress systematic false positives without enforcing a positive label for another class.

#### Spatial attention pooling

For each frame, the set of spatial tokens was reduced to a single frame-level representation using learnable spatial attention pooling. The module learned attention weights over all spatial token positions and computed a weighted sum of the token features, followed by a linear projection into a lower-dimensional feature space. The learned attention weights could be projected back into image space during inference to generate spatial attention heatmaps for interpretability. The architecture incorporated a second temporal stream computed at lower temporal resolution. Short-scale frame features were temporally upsampled to match the long-scale stream and concatenated with it, enabling the head to combine fine grained temporal detail with broader contextual information.

#### Temporal pooling and decoding

Temporal pooling over windows of fixed size averaged neighboring frame embeddings was done to reduce temporal noise. The pooled sequence was then decoded using a multi-stage temporal convolutional network (MS-TCN). An initial prediction stage produced preliminary logits, followed by one or more residual refinement stages in which each stage added a learned correction to the previous stage output. Refined logits were upsampled back to the original temporal resolution to produce framewise predictions. Clip-level logits were computed by temporal averaging of the pooled predictions.

#### Localization detection

The framework optionally supported localization supervision using framewise bounding boxes, usually around 10 bounding boxes are enough for the model to learn to track an animal. In this mode, a localization head predicted object position from spatial tokens and refined predictions temporally, enabling ROI-aware learning when tracking-like supervision was available. Note that this is different from tracking enabled by SAM2, which SBL also supports.

#### Training objectives

The total training loss was a weighted sum of modular terms, with inactive components omitted when the corresponding mode was disabled. The classification loss was the standard cross-entropy in multi-class mode or binary cross-entropy in OvR mode, with optional focal, contrastive and asymmetric loss variants for imbalance sensitive settings. Label smoothing was applied by softening binary targets away from zero and one. A hard pair mining loss enforced a margin between the predicted score of the true class and that of a confusion-prone rival class. The boundary loss used binary cross-entropy against transition indicators optionally expanded by a tolerance window around true boundaries. A temporal smoothness regularizer penalized implausible frame-to-frame fluctuations in the predicted logits, serving as a soft prior for temporally contiguous behavior bouts.

#### Sampling and augmentation

Class-aware balanced sampling was used to mitigate label imbalance, with optional confusion-aware weighting that increased sampling probability for frames and clips recently associated with high model confusion. Virtual expansion and clip-stitching augmentation were applied in low data regimes to increase training diversity without requiring additional manual labels. Data augmentations included horizontal and vertical flips, color jitter, brightness perturbation, Gaussian blur, random noise, small rotations, speed perturbation, random occluding shapes, and crop jitter in localization mode.

#### Optimization and auto-tuning of hyperparameters

All trainable components were optimized using AdamW with configurable learning rates, weight decay, scheduler based annealing, and optional exponential moving average (EMA) of weights. Because the backbone remained frozen, optimization was restricted to the compact task head. A lightweight random search auto tuning procedure was applied prior to final training. Candidate hyperparameter configurations were each trained for a shortened schedule and evaluated on validation macro F1. The best performing configuration was then used to retrain the final model from scratch with the full training budget. The selected configuration was stored as a reusable training profile.

#### Inference and postprocessing

For long videos, inference was performed using overlapping sliding windows, with framewise predictions stitched across window boundaries. Raw predictions were temporally smoothed and segmented into behavioral bouts using class-specific thresholds, minimum duration constraints, and merge-gap rules. For OvR models, per-class thresholds were calibrated on a validation set by maximizing per-class F1, and optional per-class temperature scaling was applied to improve the correspondence between raw scores and downstream decisions. Outputs included framewise class scores, behavior timelines, attention maps, overlay videos, and review candidate reports.

#### Active learning refinement

The framework supported an iterative active learning loop. After inference, clips were ranked by uncertainty for targeted review near class boundaries, and confident predictions were ranked per class for rapid enrichment of the training set. Transition mining identified candidate behavior change points using framewise predictions and boundary logits. Rejected confident predictions were stored as class-specific hard negatives. Accepted positives, reviewed transitions, and hard negatives were written back to the annotation store and reused in subsequent training iterations, forming a cycle of training, inference, review, label augmentation, and retraining.

### Behavioral state clustering

#### SAM2.1

To generate binary masks for each animal we relied on the “SAM2.1-hiera-large” model coupled with SBL to improve temporal consistency for tracking. For point prompts we developed a GUI that enables users to click on the animal (or object) of interest in the first frame, after which SBL automatically performs segmentation and tracking across frames. SAM2.1 is known to have issues with longer videos, to prevent memory issues with longer experimental recordings, videos were split into chunks of ∼200 frames and memory management optimized directly in SBL. A point prompt was automatically passed to each segment to maintain robust tracking and segmentation over the entire video.

#### Embedding extraction

Each clip was decoded frame-by-frame, resized to 288 × 288 pixels, scaled to the [0, 1] range, and passed through the pretrained VideoPrism backbone. A single D-dimensional descriptor per clip was obtained by mean-pooling over all spatiotemporal token embeddings. The resulting feature matrix was saved in compressed NPZ format alongside clip metadata (snippet ID, source group, video ID, object ID, clip index, and frame span). We implemented a metadata manager in our GUI for easier meta data management.

#### Preprocessing and dimensionality reduction

Prior to clustering, the embedding matrix was transposed into a samples × features format. Preprocessing supported four normalization modes (none, L2, z-score, and min-max); the default was no normalization, appropriate for the well-conditioned VideoPrism embedding space. For visualization, embeddings were projected into two dimensions using UMAP (n_neighbors = 30, min_dist = 0.1, random_state = 42). Clustering was performed in the full high-dimensional preprocessed space rather than on UMAP coordinates, to avoid artifacts introduced by low-dimensional projection.

#### Clustering algorithms

Two clustering algorithms were supported. Leiden community detection was applied on a k-nearest-neighbor graph (k = 15) constructed over the embedding space and partitioned using the RBConfigurationVertexPartition objective (resolution = 1.0). HDBSCAN was used as an alternative for irregular cluster geometries or when explicit noise modeling was preferred (min_cluster_size = 5, min_samples = 1, cluster_selection_epsilon = 0.0). Noise points were assigned label −1. Cluster assignments were merged back into the clip metadata table by snippet ID, enabling downstream stratification by experimental group, subject identity, or source video.

#### Representative snippet selection and interactive inspection

Representative snippets were identified per cluster by ranking member clips by Euclidean distance to the cluster centroid in the preprocessed embedding space; the ten closest clips per non-noise cluster were exported as exemplars into the labeling workflow with their cluster label as a provisional annotation. Additionally, the UMAP projection was rendered as an interactive plot in which individual points could be selected directly, launching an embedded video player displaying the corresponding clip alongside its surrounding temporal context to enable rapid assessment of behavioral states and identification.

#### Hardware and software

All experiments were conducted on a Linux workstation equipped with a single NVIDIA GeForce RTX 4080 SUPER GPU with 16 GB vRAM. Model training and inference were implemented in Python within a Conda managed environment. SBL model training used PyTorch with CUDA (12.3) acceleration. VideoPrism backbone inference was executed using JAX/Flax on the same GPU. All analyses were run on a single workstation without distributed computing.

#### AI use disclosure

Claude (Anthropic) and ChatGPT (OpenAI) were used for proofreading, language editing, and improving the readability of this manuscript. All scientific content, data interpretation, and conclusions were generated and verified by the authors alone.

## Acknowledgements

The authors would like to thank Dana Matzek and Bianca Stahr for animal husbandry and Clara de la Rosa del Val for critical input on this work. We also thank Takaaki Kaneko for providing access to their marmoset recordings for analysis with SBL.

## Author contributions

AA and FMB designed the experiments. AA, NH, FK, HP collected data. AA analyzed the data. AA and FMB wrote the manuscript. AA made the figures.

## Ethics approval

Procedures were performed in accordance with the Regierung von Oberbayern.

## Data and materials availability

All datasets used and/or analyzed in this present study are available from the corresponding author upon request.

Code repository: https://github.com/alms93/SingleBehaviorLab

## Extended Data Figures

**Extended Data Figure 1.**
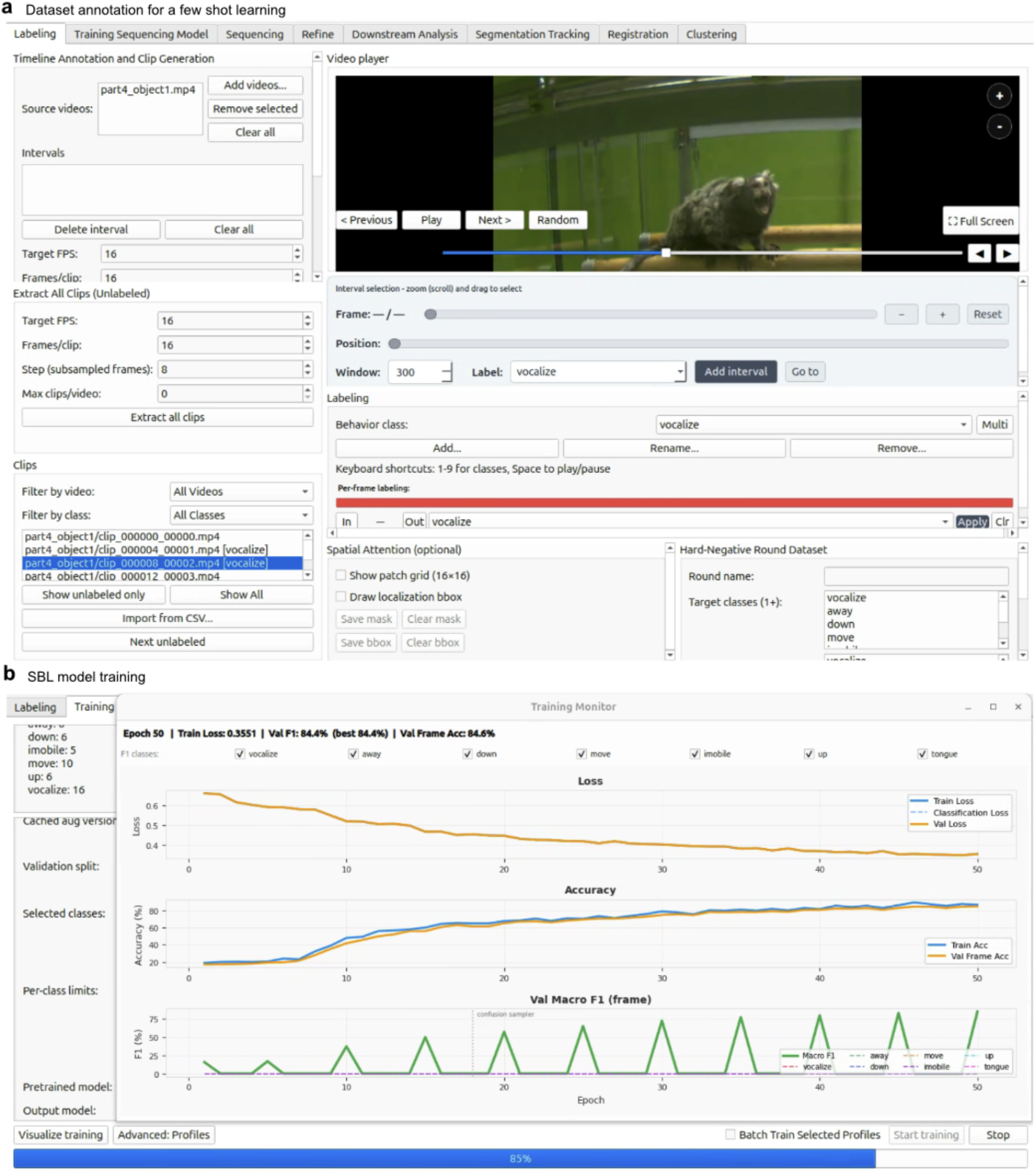
SingleBehavior Lab graphical user interface for dataset annotation and model training. **a)** Dataset annotation interface for few-shot learning. The labeling panel (left) allows users to specify source videos, define temporal intervals for clip extraction, and configure clip sampling parameters including target FPS, frames per clip, subsampled frames, and maximum clips per video. Extracted clips are organized by video and behavioral class, with options to filter by video or class and to import annotations from CSV. The video player panel (right) displays the source recording with interactive interval selection, enabling frame-precise annotation of behavioral events. Behavior classes (e.g., vocalize, away, down, move) are defined and managed directly within the interface, with support for multi-class labeling and keyboard shortcuts for rapid annotation. An optional spatial attention module allows users to define spatial regions of interest, and a hard-negative round dataset can be configured to improve classifier specificity. **b)** SBL model training interface. The training panel (left) displays configurable parameters including validation split, selected behavior classes, per-class clip limits, pretrained model selection, and output model path. The training monitor (right) displays live learning curves across epochs for training and validation loss, frame-level classification accuracy, and per-class validation macro F1 score, allowing users to assess model convergence and per-class performance in real time.

**Extended Data Figure 2.**
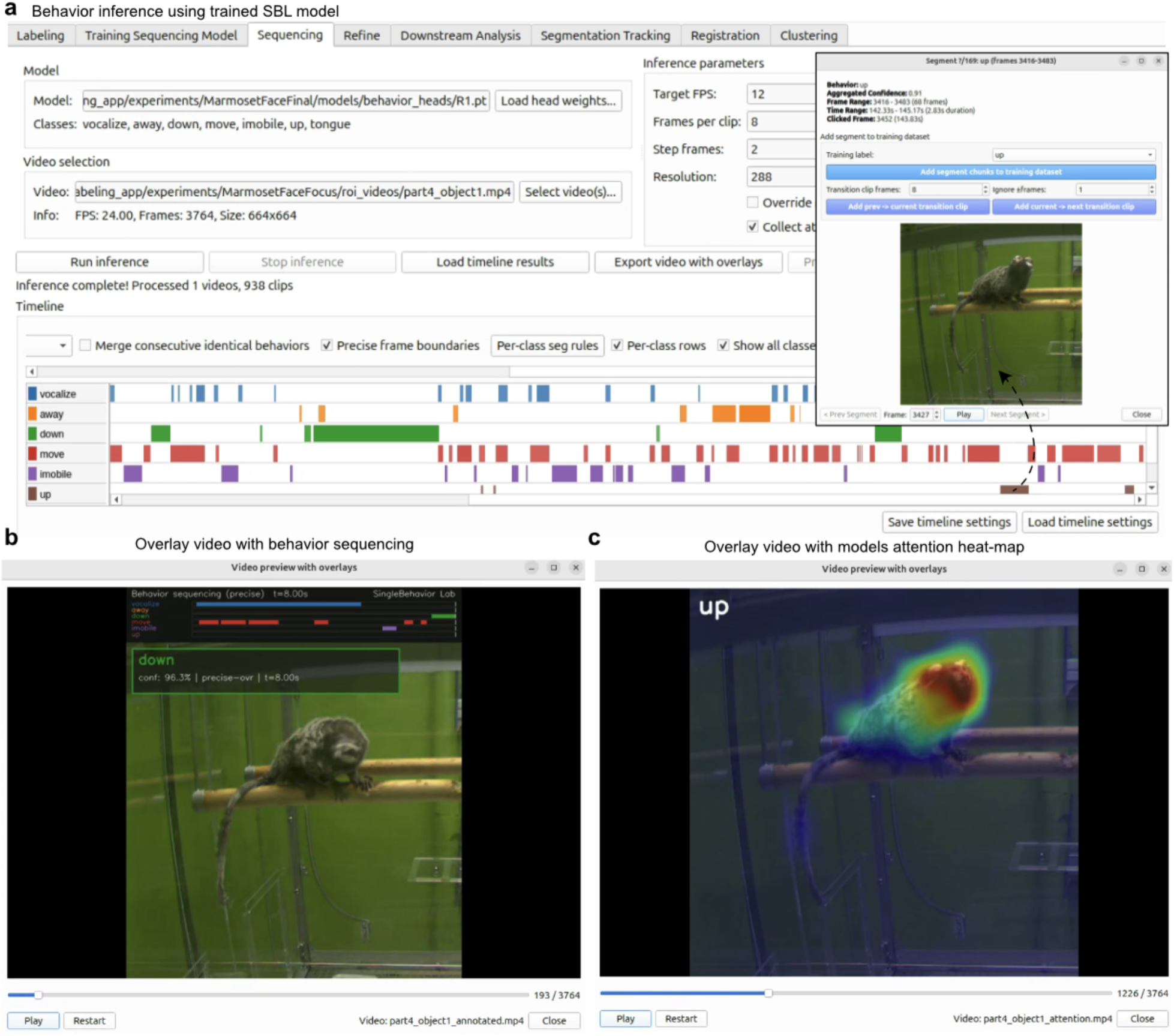
SingleBehavior Lab graphical user interface for behavior inference, timeline visualization, and model interpretability. **a)** Behavior inference interface using a trained SBL model. The model panel specifies the trained model path and the associated behavior classes (e.g., vocalize, away, down, move, immobile, up, tongue). Inference parameters including target FPS, frames per clip, step frames, and input resolution are configured prior to inference. Video selection allows specification of the source recording along with its metadata (frame count, resolution). Following inference, the interface displays a per-class behavior timeline in which each classified behavior is represented as a color-coded horizontal track across the full recording duration. Options for merging consecutive identical behaviors, enforcing precise frame boundaries, applying per-class segmentation rules, and filtering displayed classes are available directly in the timeline view. Individual timeline segments can be selected to inspect aggregated confidence scores and temporal boundaries for each classified bout. Processed timelines can be exported as annotated overlay videos or saved as timeline files for downstream analysis. **b)** Overlay video with behavior sequencing. The video preview window displays the source recording with real-time behavior annotations overlaid, including the current predicted class label (e.g., “down”), the frame-level confidence score, and the temporal extent of the active behavioral bout. A color-coded behavior timeline bar is rendered at the top of the frame, providing continuous temporal context during video playback. **c)** Overlay video with model attention heatmap. The video preview window displays the model’s spatial attention heatmap overlaid on the source recording, highlighting the image regions that most strongly contributed to the current frame-level prediction (e.g., “up”). The heatmap uses a red-to-blue color scale to indicate attention intensity, providing a frame-by-frame visualization of the spatial features driving model decisions and enabling interpretability assessment of the trained classifier.

**Extended Data Fig 3.**
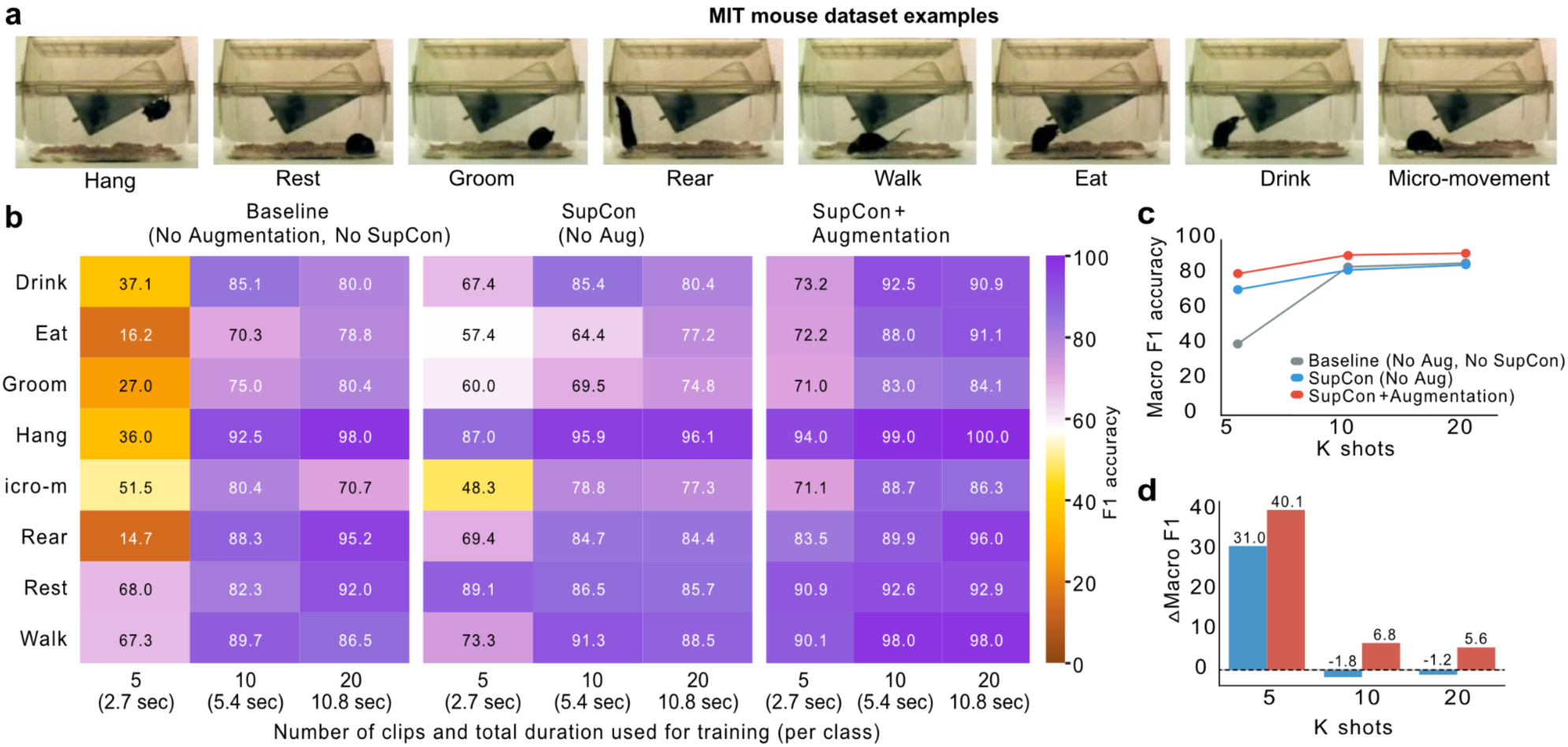
SingleBehavior Lab enables few-shot behavior classification using foundation video embeddings and a task specific contrastive adapter training. **a)** Example frames from the MIT mouse behavior dataset [33] showing the eight ground truth annotated behaviors used in this study. **b)** Heatmaps of per-class F1 accuracy for three training configurations. Baseline (no augmentation, no SupCon), SupCon without augmentation, and SupCon with augmentation. All evaluated using 5, 10, or 20 training clips per class (2.7, 5.4, and 10.8 s total duration). **c)** Macro F1 accuracy plotted as a function of the number of training clips per class for each training configuration. **d)** Difference in macro F1 accuracy relative to the baseline configuration for each training condition and number of training clips. Macro F1 represented as %.

**Extended Data Figure 4.**
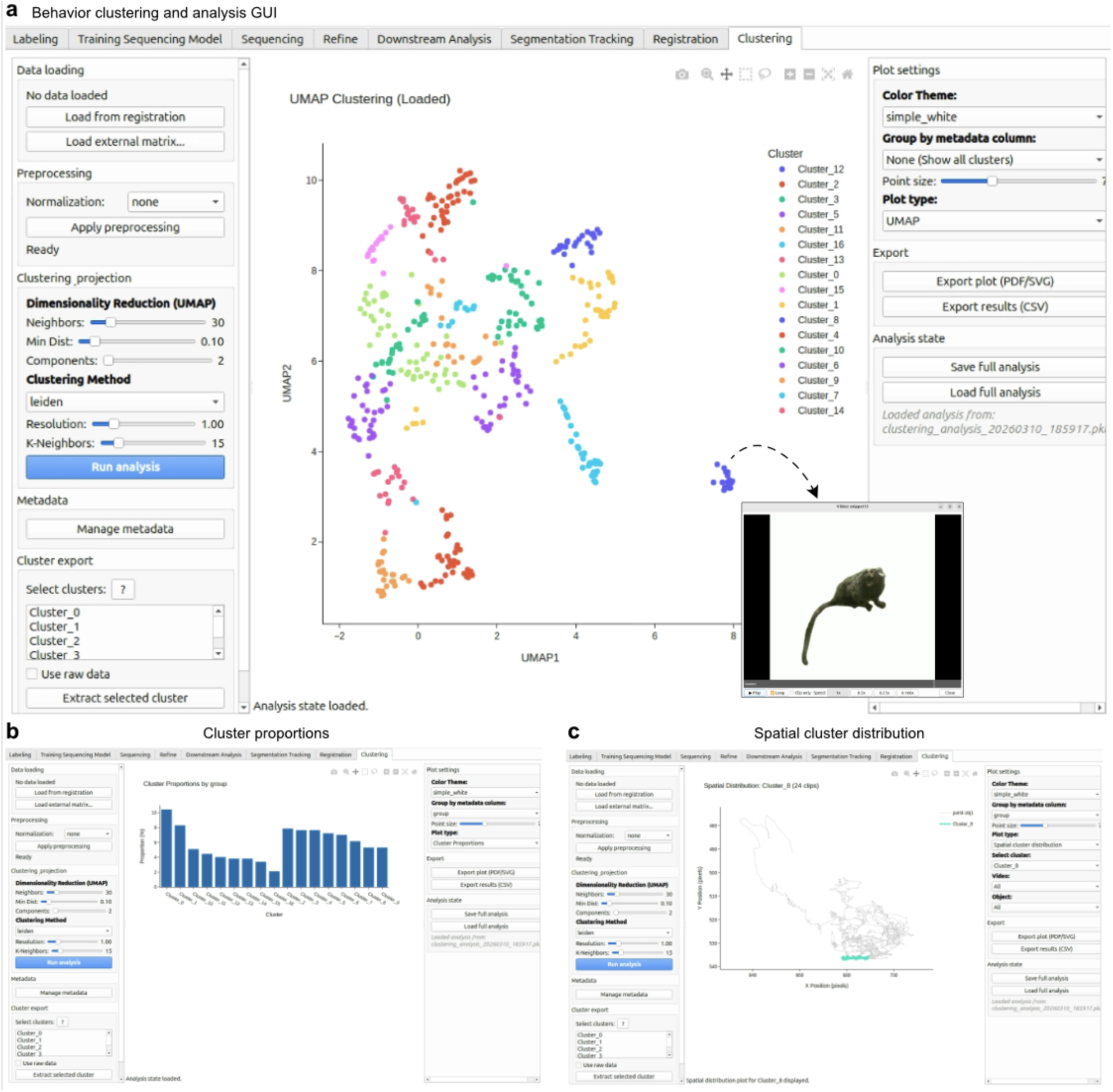
SingleBehavior Lab graphical user interface for behavior clustering and analysis. **a)** Behavior clustering interface. The left panel provides controls for data loading (from registration or external matrix), preprocessing normalization, dimensionality reduction via UMAP (configurable neighbors, minimum distance, and number of components), and clustering method selection (Leiden algorithm with adjustable resolution and k-neighbors). Following analysis, identified clusters are displayed as a UMAP scatter plot color-coded by cluster identity. Individual video clips corresponding to selected data points can be previewed directly within the interface. Cluster assignments and full analysis states can be exported as CSV or saved for reloading. **b)** Cluster proportions view. Displays the relative abundance of each behavioral cluster across experimental groups as a bar chart, enabling direct quantitative comparison of cluster composition between conditions. **c)** Spatial cluster distribution view. Maps the spatial distribution of behavioral clusters onto the animal’s position within the recording arena, allowing identification of behaviors associated with specific spatial locations.

**Extended Data Figure 5.**
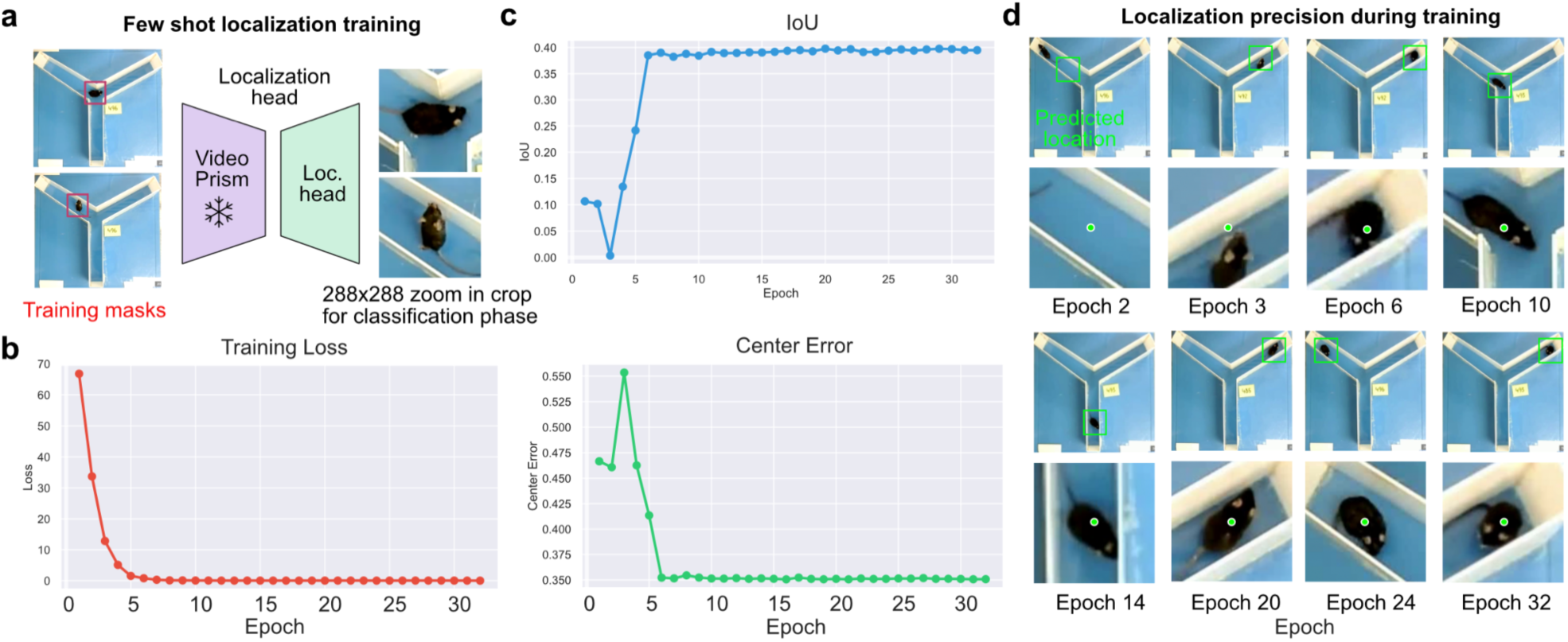
Few-shot localization training and convergence of the SBL localization head. **a)** Schematic of the few-shot localization training pipeline. Annotated training masks are used to supervise a lightweight localization head trained on top of frozen VideoPrism embeddings. The localization head predicts animal position, and the resulting 288×288 pixel crop is used as input to the classification phase. **b)** Training loss and center error across epochs. Training loss decreases rapidly within the first five epochs and plateaus at low values, while center error follows a similar trajectory, indicating fast convergence of the localization head with minimal supervision. **c)** Intersection over Union (IoU) across training epochs. IoU increases steeply in early epochs and stabilizes above 0.90, confirming that the localization head learns accurate and spatially precise animal crops within a small number of training steps. **d)** Qualitative localization precision across epochs. Representative frames at selected epochs (2, 3, 6, 10, 14, 20, 24, 32) illustrate progressive improvement in predicted crop placement, with accurate centering on the animal achieved by approximately epoch 10 and maintained thereafter.

**Extended Data Fig. 6.**
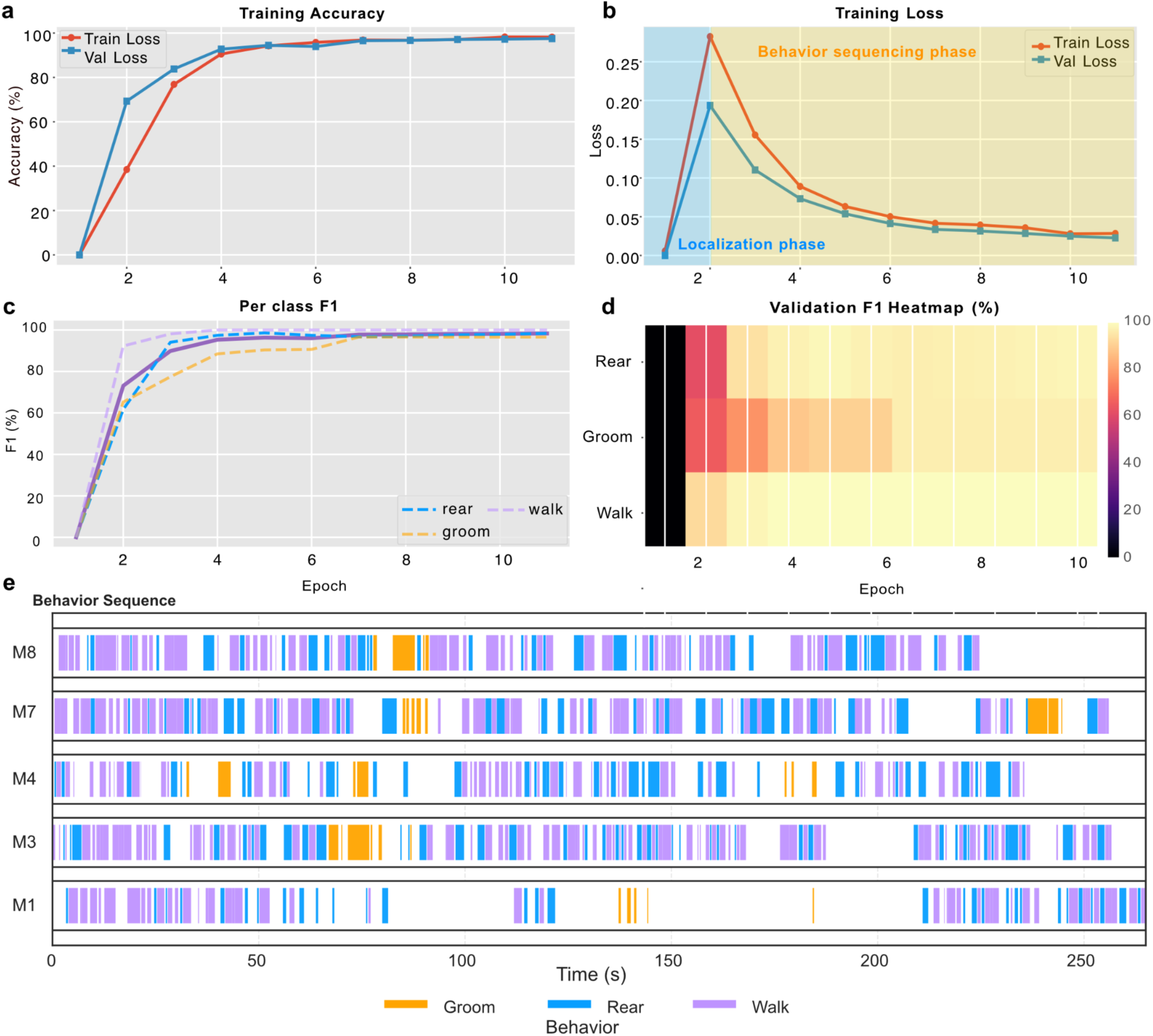
Training dynamics and class-wise performance of the few-shot behavior sequencer. **a)** Training and validation accuracy across epochs (red, training; blue, validation). Both metrics increase rapidly within the first two epochs and plateau above 95%. **b)** Training and validation loss curves (red, training; teal, validation), with two annotated training phases, an initial localization phase (blue shading) and a subsequent behavior sequencing phase (yellow shading). Loss peaks transiently at the phase transition before decreasing steadily across both splits. **c)** Per-class F1 scores across training epochs (blue dashed, rear; orange dashed, groom; purple dashed, walk). All classes stabilize above 95% F1 by epoch 6, demonstrating balanced classification across behavioral categories. **d)** Validation F1 heatmap (%) across training epochs for each behavioral class, showing progressive improvement converging to high F1 values by mid-training. **e)** Predicted behavior sequences for five representative mice (M1, M3, M4, M7, M8) (orange, groom; blue, rear; purple, walk).

**Extended Data Fig. 7.**
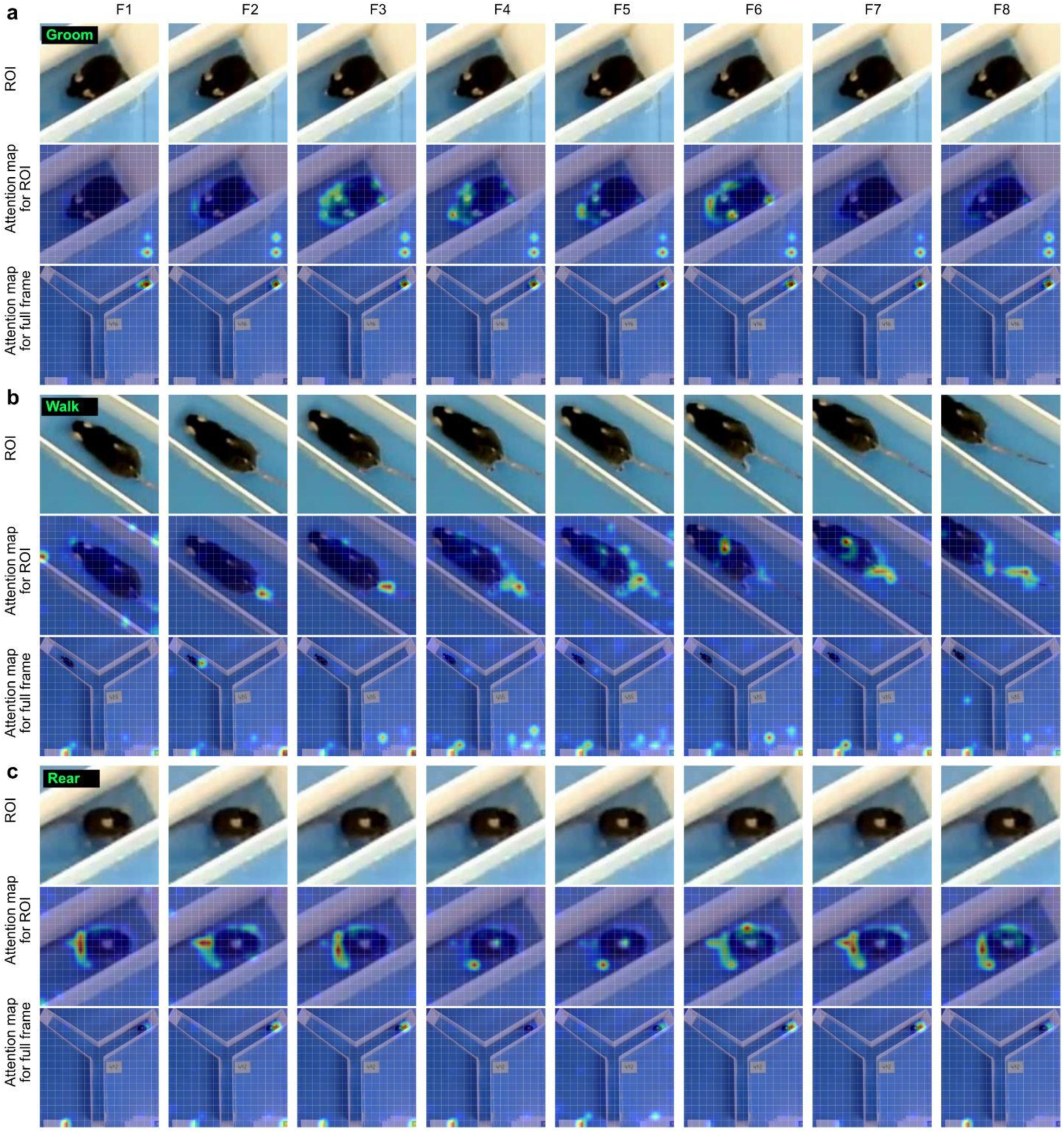
Attention maps reveal behavior-specific spatial features learned during behavior sequencing. a–c. Representative sequences (F1–F8) illustrating model attention during classification of grooming (**a**), walking (**b**) andrearing (**c**) . For each behavior, the top row shows the region of interest (ROI) input frames, the middle row shows attention maps computed over the ROI, and the bottom row shows attention maps projected onto the full-frame input.

## Additional Supplementary Data

**Supplementary Video 1. Demo video of behavior sequencing in a freely moving mouse in an open cage. <u>Suplementary_Video_1.mp4</u>**

**Supplementary Video 2. Y maze mouse localization and behavior sequencing. <u>Suplementary_Video_2.mp4</u>**

**Supplementary Video 3. Behavior sequencing for individual fly. <u>Suplementary_Video_3.mp4</u>**

